# PIEZO-dependent mechano-sensing of the niche is essential for intestinal stem cell fate decision and maintenance

**DOI:** 10.1101/2024.09.06.611641

**Authors:** Meryem B. Baghdadi, Ronja M. Houtekamer, Louisiane Perrin, Abilasha Rao-Bhatia, Myles Whelen, Linda Decker, Martin Bergert, Carlos Pérez-Gonzàlez, Réda Bouras, Giacomo Gropplero, Adrian KH Loe, Amin Afkhami-Poostchi, Xin Chen, Xi Huang, Stephanie Descroix, Jeffrey L. Wrana, Alba Diz-Muñoz, Martijn Gloerich, Arshad Ayyaz, Danijela Matic Vignjevic, Tae-Hee Kim

## Abstract

Stem cells continuously perceive and respond to various environmental signals to maintain homeostasis. In addition to biochemical factors, the stem cell niche is subjected to mechanical and physical cues. However, it remains unclear how stem cells can sense mechanical signals from their niche *in vivo*. Since intestinal stem cells constantly and directly face the external environment, we investigated the roles of mechano-sensing PIEZO ion channels in the gut stem cell niche. By employing mouse genetics and performing single-cell RNAseq analysis, we revealed the absolute requirement for PIEZO channels in intestinal stem cell (ISC) state dynamics and maintenance. *In vivo* measurement of basement membrane region stiffness demonstrated that ISCs reside in a more rigid microenvironment at the bottom of the crypt. Using 3D and 2D organoid systems combined with bioengineered substrates and a cell stretching device, we found that PIEZO channels are activated by high extracellular matrix stiffness and tissue tension to modulate ISC behavior. This study delineates the mechanistic cascade of PIEZO channel activation in ISCs from the upstream extracellular stimuli through the downstream signaling activation that coordinates stem cell fate decision and maintenance.

## INTRODUCTION

Tissue turnover and regeneration are orchestrated by stem cells that both differentiate and self-renew. The balance of self-renewal, proliferation, and commitment is tightly regulated by the stem cell microenvironment, or “niche” (*1*, *2*). It is thus essential to understand how stem cells integrate niche signals to maintain homeostasis. The intestinal epithelium constitutes an excellent paradigm for studying stem cell biology, as it withstands continuous renewal and has a striking ability to regenerate upon acute and chronic injury (*3*). LGR5+ intestinal stem cells (ISCs) located at the bottom of the crypt give rise to transit-amplifying cells (TAs) that commit to secretory or absorptive progenitors while migrating up in the crypt. The stem cell-specific microenvironment provides key paracrine factors (e.g., WNT, NOTCH, BMP, etc.) that tightly regulate stem cell proliferation, self-renewal, and differentiation (*3*, *4*).

In addition to these biochemical cues, crypts are also characterized by a specific geometry, an extracellular matrix (ECM) composition and distribution that may contribute to local stiffness and tissue tension (*5*). In addition, ISC proliferation induces mechanical stress on neighboring cells at the bottom of the crypt (*6*). These mechanical cues might impact ISC behavior, but how intestinal stem cells integrate the mechanical signals from their niche *in vivo* remains unclear. It is also unknown if mechano-sensing is required for stem cell maintenance.

Cells sense and respond to mechanical cues via the integrated activation of different signaling pathways. This mechanotransduction process eventually leads to changes in cell shape, gene expression, and cell fate (*7*). PIEZO ion channels, which include PIEZO1 and PIEZO2, are an important group of mechanosensitive signaling receptors. These channels open in response to mechanical stimuli, allowing calcium to flow into the cell, activating downstream signaling pathways (*8–10*). PIEZO channels play major roles in various aspects of mechanotransduction in mammals, and the mutations in these channels are associated with several human diseases (*11*, *12*). Specifically, it has been shown that PIEZO1 activation, triggered by substrate stiffness, directs lineage decisions in human neural stem cells *in vitro* (*13*). In addition, when mechanically activated, *Drosophila* PIEZO, which has ∼24% sequence identity with mammalian PIEZO channels and is specifically expressed in enteroendocrine precursor cells, triggers cell differentiation. However, the loss of *Piezo* induces no changes in stem cell numbers, indicating they are dispensable for stem cell maintenance (*14*). As the *Drosophila* gut lacks the typical crypt-villus structure, implying distinct mechanical niche properties, the *in vivo* roles of mammalian PIEZO channels in intestinal stem cell activity remain to be determined.

To define the specific roles of PIEZO channels in ISCs, we performed *in vivo* conditional genetic ablation of *Piezo* genes in the intestinal epithelium. Strikingly, *Piezo 1* and *2* double knockout (KO) mice exhibited stem cell depletion, leading to lethality. Through single-cell transcriptome analysis of *Piezo* knockout mice, we identified NOTCH and WNT as downstream pathways involved in PIEZO signal transduction. I*n vivo* AFM measurement of the basement membrane region stiffness confirmed that ISCs experienced a more rigid microenvironment at the bottom of the crypt compared to cells residing at the top. Using 3D and 2D organoids combined with bioengineered substrates of varying rigidities and a tissue stretcher, we revealed that PIEZO channels activated by high substrate stiffness and tissue tension modulate ISC behavior.

## RESULTS

### PIEZO mechanosensitive channels are essential for ISC function

To define the expression pattern of PIEZO channels in intestinal epithelial cells, we analyzed an RNAseq dataset of FACS-purified cell populations isolated from mouse colon (*15*) and found that *Piezo1* is expressed in all epithelial cell types, including stem cells, while *Piezo2* is only expressed in enterocytes and enteroendocrine cells (**Fig. 1A**). We confirmed these observations by smFISH staining showing that *Piezo1* is highly expressed in most crypt cells whereas *Piezo2* expression was low (**Fig. 1B and Fig. S1A).** At the protein level, PIEZO1 was mostly detected at two specific locations: 1) at the top of the villus as cytoplasmic protein aggregates in extruding cells (**Fig. 1C**) which consistent with its known role in cell extrusion and 2) the crypt cells, enriched at the basolateral membrane (**Fig. 1C**).

**FIGURE 1:**
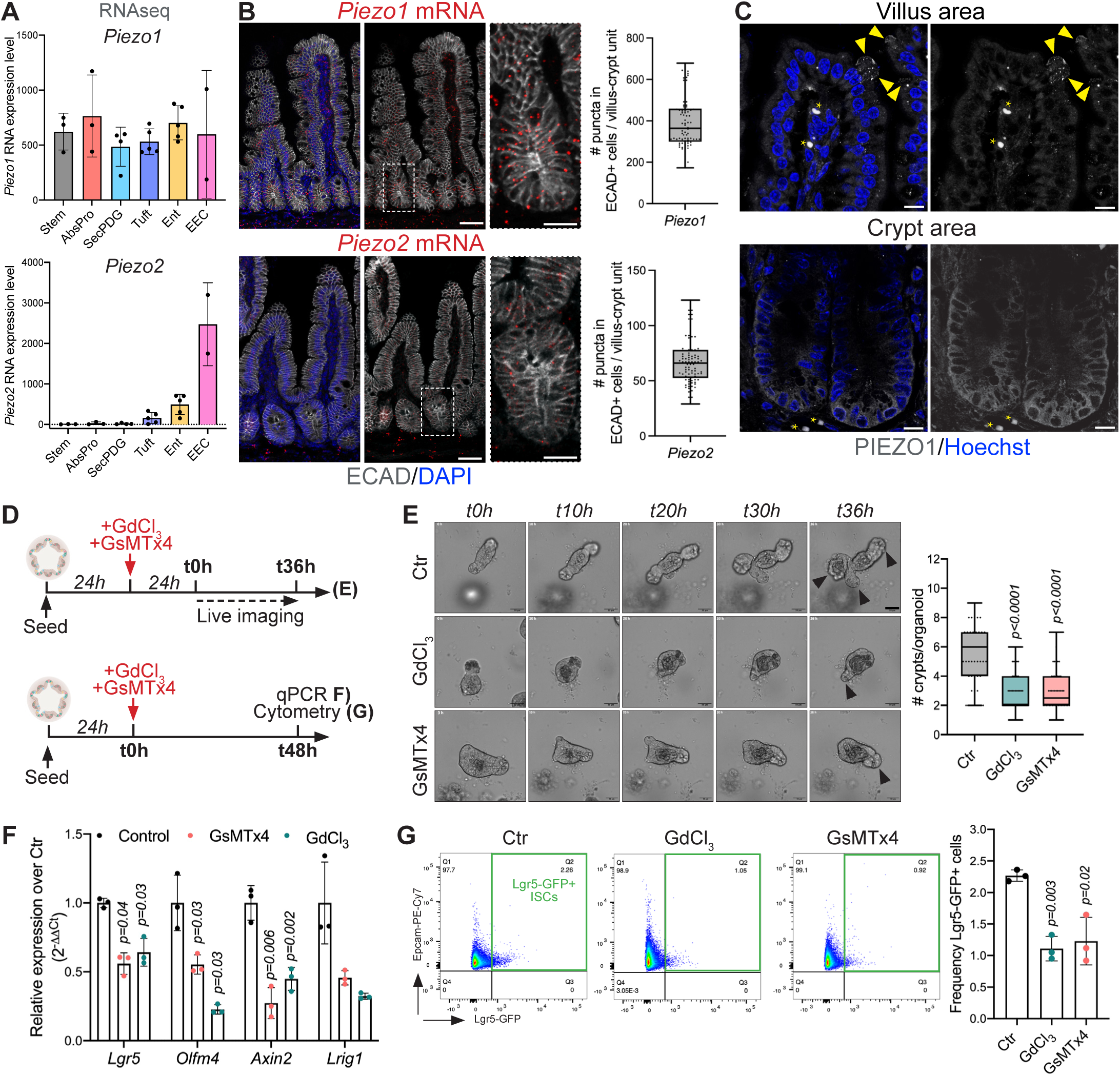
PIEZO channel inhibition impairs ISC marker gene expression and numbers. **(A)** *Piezo1* and *Piezo2* gene expression levels in FACS-sorted colonic epithelial cells assessed by RNA deep sequencing (GEO accession number GSE143915). FACS isolated six distinct populations including stem, absorptive progenitor (AbsPro), secretory progenitors/deep crypt secretory cells/goblet cells (SecPDG), tuft cells, enterocytes (Ent), and enteroendocrine cells (EEC). **(B)** smFISH for *Piezo1* (left) and *Piezo2* (right) expression combined with E-Cadherin (ECAD) immunostaining to visualize intestinal epithelial cells and DAPI (DNA) (n=90 crypts from N=4 mice). Insets show signal specifically in the crypts. Scale bar, 50μm and 10μm in insets. **(C)** PIEZO immunostaining on WT intestines showing PIEZO signals at the tip of the villus (arrowheads label extruding cells) and at the bottom of the crypt. Asterisks show positive red blood cells. Scale bar, 10μm. **(D)** Experimental scheme of *in vitro* PIEZO channel pharmacological inhibition in *Lgr5-GFP* intestinal organoids, using GdCl_3_ or GsMTx4. (Top) Live imaging was performed for 36h with a timeframe of 1h. (Bottom) qRT-PCR and flow cytometry analysis was performed after 48h of PIEZO inhibition. **(E)** Representative images of brightfield time-lapse imaging of control (Ctr), GdCl_3_, and GsMTx4-treated organoids. See Movies S1 to S3. Quantification of *de novo* crypts formed per organoid (n=100 organoids from n=3 independent experiments). Scale bar, 50μm. **(F)** Transcript analysis of stem cell markers in control organoids (Ctr) or organoids treated with PIEZO inhibitors (GdCl_3_ or GsMTx4) (n=3 independent experiments). **(G)** Representative gating and quantification of ISC frequency by flow cytometry in control organoids (Ctr) or organoids treated with PIEZO inhibitors (GdCl_3_ or GsMTx4) (n=3 independent experiments). Error bars indicate SD. Kolmogorov-Smirnov test (B, E), Two-tailed paired Student’s *t*-test (F, G), and *P* values are shown in panels. Box edges show 25th and 75th percentiles, central point is median, and error bars represent minimum and maximum values. See also Supplementary Figure S1.

To assess whether PIEZO channels regulate stem cell function, we blocked their activation using two pharmacological inhibitors (GdCl3 and GsMTx4; (*8*, *9*, *16*)) in ISC-containing 3D organoids (**Fig. 1D**). Live imaging revealed that PIEZO inhibition impaired organoid growth and, the number of *de novo* crypts per organoid (**Fig. 1E-F** and **Movies S1-S3**). Consistently, PIEZO inhibition decreased the expression of stem cell markers (**Fig. 1D and 1F**) and the frequency of Lgr5-GFP^+^ stem cells per organoid (**Fig. 1G**). Together, these results suggest that PIEZO channels are required for maintaining ISC numbers.

To examine PIEZO function in gut homeostasis *in vivo*, we crossed *Villin-creERT2* transgenic mice with either *Piezo1^Flox^* or *Piezo2^Flox^* to generate *Piezo1^cKO^* and *Piezo2^cKO^* mice in which the expression of *Piezo1* or *Piezo2* was ablated from all intestinal epithelial cells, including stem cells upon tamoxifen administration (**Fig. S2A-C**). These mice were healthy, showed no morphological nor histological intestinal defects (**Fig. S2D-E**) and harbored typical ISC number (**Fig. S2F**). Interestingly, we found that in *Piezo1*-ablated mice (*Piezo1^cKO^*), the *Piezo2* expression level was upregulated 4-fold (**Fig. S2G**), suggesting that *Piezo2* could compensate for the lack of *Piezo1* to maintain intestinal stem cell homeostasis.

To test the potential redundancy between PIEZO1 and PIEZO2 channels, we generated double knockout mice in which both PIEZO channels were absent from intestinal epithelial cells upon tamoxifen induction (*Villin-creERT2;Piezo1^Flox^*;*Piezo2^Flox^*; herein *Piezo^dbKO^*) (**Fig. 2A and Fig. S3A-C**). Strikingly, six days post tamoxifen treatment, *Piezo^dbKO^* mice exhibited diarrhea, blood in stool, and rapid decline of body weight (loss of >20% of the initial body weight), leading to lethality. Moreover, histological analysis of *Piezo^dbKO^* mutant intestines showed longer crypts (**Fig. 2B**) and shorter villi (**Fig. 2C**). Thus, these data revealed that PIEZO channels in intestinal epithelia are essential for maintaining adequate intestinal architecture and homeostasis.

**FIGURE 2:**
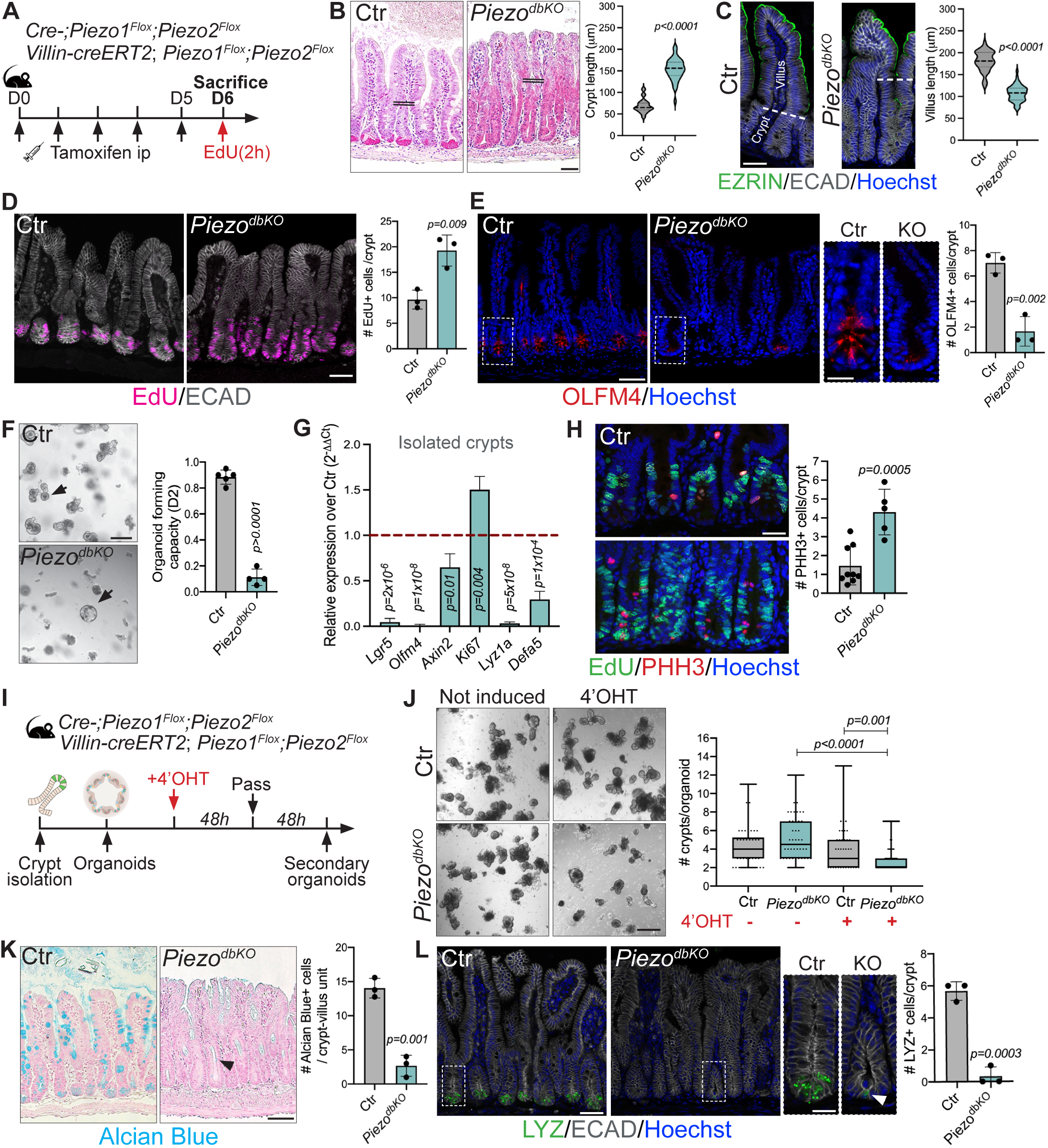
Genetic ablation of PIEZO channels induces stem and secretory cell depletion. **(A)** Experimental scheme of control (Ctr; *Cre-;Piezo1^Flox^;Piezo2^Flox^*) and *Piezo1* and *Piezo2* conditional double knock-out mice (*Piezo^dbKO^*; *Villin-creERT2;Piezo1^Flox^;Piezo2^Flox^*). Mice were injected intraperitoneally (ip) with tamoxifen once a day for 5 consecutive days and sacrificed at day 6 (D6). EdU was injected 2h before sacrifice. **(B)** Representative images of Hematoxylin/Eosin staining of intestinal transverse sections from control (Ctr) and *Piezo* double KO mice (*Piezo^dbKO^*). Quantification of crypt length (n=40 crypts from N=3 mice). Double black lines represent crypt-villus boundary. Scale bar, 25μm. **(C)** EZRIN immunostaining showing the brush border and quantification of villus length (n=50 villus from N=3 mice). Dashed white lines showing the crypt-villus boundary. ECAD immunostaining (white) was used to visualize epithelial cells. Scale bar, 25μm. **(D)** EdU labeling and quantification from control (Ctr) and *Piezo* double KO mice (*Piezo^dbKO^*) (n=150 crypts from N=3 mice). Scale bar, 50μm. **(E)** Immunostaining for OLFM4 and quantification from Ctr and *Piezo^dbKO^*mice (n=70 crypts from N=3 mice). Scale bar, 50μm and 10μm in inset. **(F)** Organoid formation from crypts isolated from Ctr and *Piezo^dbKO^* mice. Quantification of the organoid forming capacity (= number of organoids formed/number of plated crypts) at day 2 of culture (N=4 mice). Scale bar, 250μm. **(G)** qRT-PCR analysis of stem cell (*Lgr5, Olfm4, Axin2*), proliferation (*Ki67*) and Paneth cell (*Lyz1a, Defa5*) markers in crypts isolated from Ctr and *Piezo^dbKO^* mice (N=3 mice). Red line represents gene expression in the Ctr (=1). **(H)** Phospho-histone 3 (PHH3) immunostaining and EdU labeling and quantification in Ctr and *Piezo^dbKO^* mice (n=50 crypts from N=3 mice). Scale bar, 50μm. **(I)** Experimental scheme of *in vitro Piezo* genetic ablation in organoids using 4-hydroxytamoxifen (4’OHT) treatment for 48h. **(J)** Self-renewal capacity of secondary organoids (3 days upon sub-culturing) isolated from Ctr and *Piezo^dbKO^* mice and induced with 4-hydroxytamoxifen (4’OHT). Non-induced organoids were used as controls (n=60 organoids from N=3 mice). Scale bar, 250μm. **(K)** Alcian Blue staining of secretory cells and quantification from Ctr and *Piezo^dbKO^* mice (n=150 crypts from N=3 mice). The arrow points to a residual secretory cell. Scale bar, 100μm. **(L)** Immunostaining and quantification of Paneth cells (LYSOZYME; LYZ+) from Ctr and *Piezo^dbKO^* (n=100 crypts from N=3 mice). The arrow in the inset shows residual Paneth cells. Scale bar, 50μm and 10μm in inset. Error bars indicate standard deviation (SD). Kolmogorov-Smirnov test (B-C), Two-tailed unpaired Student’s *t*-test (D-H, K, L), Mann-Whitney test (J). Box edges show 25th and 75th percentiles, central point is median, and error bars represent minimum and maximum values. *P* values are shown in panels. See also Supplementary Figure S3.

Consistent with crypt hyperplasia, *Piezo^dbKO^* mice showed a significant increase of 5-ethynyl-20-deoxyuridine (EdU) incorporation, reflecting ectopic epithelial cell proliferation (**Fig. 2D**). Importantly, PIEZO ablation induced OLFM4+ ISC depletion (**Fig. 2E**), with a dramatic decrease of *Lgr5* expression (**Fig. S3D**), demonstrating that PIEZO channels are required for ISC maintenance *in vivo*. To further analyze ISCs at the functional level, we isolated crypts from control (Ctr) and *Piezo^dbKO^* mice and grew them as 3D organoids. Interestingly, purified crypts from *Piezo*-ablated mice did not form 3D organoid structures (**Fig. 2F**), which can be explained by the decline of ISC markers (*Lgr5*, *Olfm4* and *Axin2*; **Fig. 2G and Fig. S3D**). Of note, *Piezo^dbKO^*cells migrate efficiently along the crypt-villus axis (**Fig. S3E**). Additionally, PIEZO1 is important for cell cycle progression (*17*, *18*), thus, the increase of EdU+ cells (**Fig. 2D**) could be due to an inability to progress through cell cycle. Labeling of cells in M-phase with Phospho-histone3 marker (PHH3) showed an increase in mitotic PHH3^+^ cells upon *Piezo* deletion (**Fig. 2H**) suggesting that KO cells are unlikely blocked in S-phase. We then assessed apoptosis using TUNEL labeling and did not find any apoptotic+ cell in the crypts upon PIEZO ablation (**Fig. S3F**), suggesting that ISC loss might be driven by self-renewal and/or differentiation defects rather than cell death or cell cycle progression defects. First, we assessed whether PIEZO KO could affect ISC self-renewal capacity by subculturing 3D crypt organoids purified from non-induced *Piezo^dbKO^* mice (**Fig. 2I**). In this model, PIEZO deletion was achieved by 4-hydroxytamoxifen (4’OHT) treatment *in vitro* after organoids were formed (**Fig. 2I** and **Fig. S3G** for efficiency of recombination), and organoids were analyzed after passaging. Interestingly, after only one subculturing step, *Piezo*-depleted organoids showed a significant decrease in *de novo* crypt formation (**Fig. 2J**), representing a severe self-renewal defect. Together, these results demonstrate that PIEZO channels are required for ISC self-renewal and maintenance.

Interestingly, *Piezo* KO organoids induced with 4’OHT also showed a decrease in *Lgr5* expression (**Fig. S3H**), which is consistent with the *in vivo Piezo^dbKO^* phenotype (**Fig. S3C** and **Fig. 2C**). In addition, while the expression of the absorptive marker *Fabp* remained unchanged, the secretory cell marker *Tff3* was dramatically decreased in the absence of PIEZO (**Fig. S3H**), suggesting a defect in cell fate decision. To assess stem cell differentiation function *in vivo*, we then stained secretory cells using Alcian Blue (**Fig. 2K**) and the enterocyte brush border using EZRIN immunolabeling (**Fig. 2C**). Intriguingly, while absorptive cells were still present (**Fig. 2C**), *Piezo^dbKO^* mice exhibited a dramatic loss of secretory cells (**Fig. 2K**). Consistently, ablation of PIEZO channels induced complete loss of Paneth cells (**Fig. 2L**). These data reveal that PIEZO channels are required for stem cell differentiation into the secretory lineage.

### Activation of PIEZO channels regulates NOTCH and WNT downstream pathways

We previously established that 6 days post tamoxifen induction, *Piezo*-ablated mice exhibited stem cell loss, hyperproliferation, and differentiation defects (**Fig. 2**). To assess how PIEZO channel ablation affects intestinal crypt cell heterogeneity, we performed single-cell RNA sequencing (scRNAseq) of intestinal epithelial cells isolated from control and *Piezo^dbKO^* mutant mice 4 days after tamoxifen induction (**Fig. 3A**). By analyzing crypt cell populations 2 days earlier, we aimed to understand the initial events of ISC dynamics immediately upon PIEZO loss. Unsupervised graph clustering partitioned crypt cells into 13 epithelial populations that we annotated using previously established markers (*19–21*) (**Fig. 3B-C** and **Fig. S4A**). Interestingly, although cell number in the stem cell cluster 1 was slightly increased in *Piezo^dbKO^*mutant mice (1.4-fold; **Fig. 3C**), the number of cells expressing ISC markers (*Lgr5*, *Olfm4*, *Ascl2*) was significantly downregulated in this population (**Fig. 3D** and **Fig. S4B-D**). On the contrary, the number of cells expressing proliferation markers (*Ki67*, *Pcna*, *Mcmc5*, *Mcm6*) was highly upregulated in *Piezo^dbKO^* stem cell population (cluster 1) (**Fig. 3E** and **Fig. S4C-D**). These data suggest that upon *Piezo* deletion, ISCs lose their stemness and acquire a highly proliferative phenotype that will eventually lead to their depletion.

**FIGURE 3:**
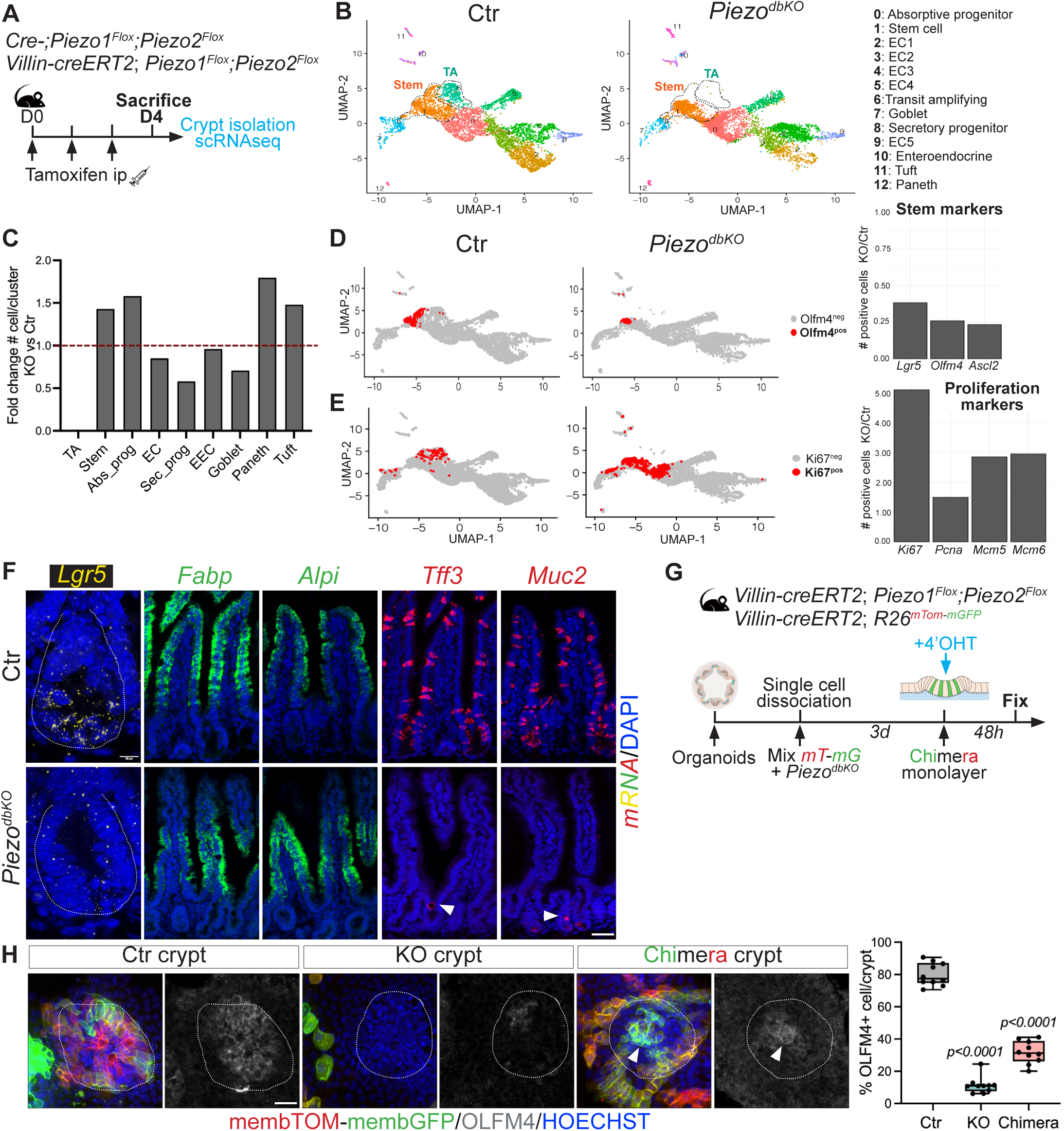
PIEZO ablation impairs stem cell state dynamics and secretory lineage specification. **(A)** Experimental scheme of single cell RNA sequencing of intestinal crypts isolated from control (Ctr; *Cre-;Piezo1^Flox^;Piezo2^Flox^*) and *Piezo1* and *Piezo2* conditional double knock-out mice (*Piezo^dbKO^*; *Villin-creERT2;Piezo1^Flox^;Piezo2^Flox^*). Mice were injected intraperitoneally (ip) with tamoxifen once a day for 3 consecutive days and sacrificed on day 4 (D4). **(B)** Uniform Manifold Approximation and Projection (UMAP) map of intestinal mouse crypt epithelial cells isolated from control (Ctr; 4713 cells) and *Piezo* double KO mice (*Piezo^dbKO^*; 6265 cells). Descriptive cluster labels are shown. EC: Enterocytes. **(C)** Fold change number of cells per cluster over control crypts. Red line represents no change (=1). **(D)** UMAP plot of *Olfm4* expression in Ctr and *Piezo^dbKO^* mice. Fold change number of cells expressing stem cell markers (*Lgr5, Olfm4, Ascl2*) in *Piezo^dbKO^* over Ctr. **(E)** UMAP plot of *Ki67* expression in Ctr and *Piezo^dbKO^* mice. Fold change number of cells expressing proliferation markers (*Ki67, Pcna, Mcm5, Mcm6*) in *Piezo^dbKO^*over Ctr. **(F)** smFISH of *Lgr5* (stem cell marker), *Fabp*, *Alpi* (absorptive markers) and *Tff3* and *Muc2* (secretory markers) in Ctr and *Piezo^dbKO^* mice (N=4 mice). Arrowheads point at residual secretory cells. Scale bar, 50μm. **(G)** Experimental scheme of chimeric 2D organoid monolayer: control (*Villin-creERT2/+;R26^mTmG^*) and *Piezo^dbKO^* organoids were dissociated into single cells, mixed together at 1:1 ratio, and plated on PAA gels to form monolayers. After 3 days in culture, monolayers were treated with 4’OHT to induce *Piezo* deletion. **(H)** Immunostaining of membrane-tomato, membrane-GFP (from control cells) and OLFM4 in chimeric monolayers. Green and/or red crypts are labelled as Ctr, “black” crypts are deleted for *Piezo* and mosaic crypts include both cell types. Quantification of percentage of OLFM4+ cells per crypt (n=10 crypts type from N=2 monolayers). Dashed line delineates crypt region and arrowhead point at OLFM4+ cells only present in PIEZO-WT cells. Error bars indicate standard deviation (SD). Mann-Whitney test (H). Box edges show 25th and 75th percentiles, central point is median, and error bars represent minimum and maximum values. *P* values are shown in panels. See also Supplementary Figure S4.

Moreover, the uncommitted multipotent transit amplifying precursor (TA) population (cluster 6) was completely absent in *Piezo^dbKO^*mutant mice (**Fig. 3B-C**). In contrast, the absorptive progenitor cluster 0 was amplified 1.6-fold (**Fig. 3C**) while the secretory progenitor cell, cluster 8, was reduced by 50% (**Fig. 3C**), indicating that multipotent TA cells principally committed towards absorptive lineage. To confirm this lineage bias *in vivo,* we performed smFISH of lineage markers at day 4 post-induction. Although *Alpi* and *Fabp* (absorptive markers) expression remained unchanged, *Tff3* and *Muc2* secretory markers were lost upon *Piezo* deletion (**Fig. 3F**). Altogether, these results demonstrate that in the absence of PIEZO, ISC self-renewal ability decreases dramatically while they adopt a highly proliferative unipotent transit amplifying identity, leading to stem cell depletion (**Fig. S4D**). In addition, pre-existing multipotent TA cells differentiate preferentially into absorptive progenitors, leading to secretory cell deficiency in the intestine (**Fig. S4D**).

Paneth cells are important stem cell regulators, providing both NOTCH and WNT ligands (*22*). Interestingly, at day 4 post-induction, while ISCs were already lost, Paneth cells were still present (**Fig. 3C** and **Fig. S4D**) and showed no changes in their NOTCH ligand expression (**Fig. S5A**). Although *Wnt3a* expression was decreased in mutant Paneth cells (**Fig. S5A**), this alone is unlikely to explain the complete loss of ISCs, as WNT ligands are also provided by the mesenchymal niche (*4*).

To test whether *Piezo* influences ISCs intrinsically, we mixed dissociated cells from control (*VillinCre-ERT2/+;R26^mTmG^*) and *Piezo^dbKO^* 3D organoids and seeded them on collagen I and laminin1-coated polyacrylamide (PAA) gels as monolayers. This model recapitulates the compartmentalization of the main cell types of the intestinal epithelium with distinguishable crypt and villus domains (*23*). While the PIEZO-WT crypts contained OLFM4+ ISCs and the PIEZO-KO crypts did not, the mixed crypts had reduced number of OLFM4+ cells (**Fig. 3H**). More specifically, within the same mosaic crypt, OLFM4+ cells were found only in PIEZO-WT cells (**Fig. 3H**). Importantly, these results demonstrated that PIEZO-WT cells were not able to rescue the phenotype of the neighboring PIEZO-KO cells suggesting that PIEZO mechanotransduction regulate ISC function in a cell intrinsic manner.

As PIEZO channels convert a mechanical input into an intracellular biological signal, we investigated PIEZO downstream pathways implicated in ISC regulation. First, we examined NOTCH signaling as it promotes crypt proliferation (through *Hes1* activation) and inhibits secretory lineage differentiation (through *Atoh1* inhibition) (*24–27*). Notably, the phenotype of *Piezo^dbKO^* mice was reminiscent of the constitutive activation of NOTCH signaling in the mouse intestinal epithelium. Indeed, NOTCH-overexpressing mice also showed an expansion of the proliferative zone and repression of secretory cell differentiation (*25*, *28*, *29*). To assess whether NOTCH signaling was altered upon *Piezo* deletion, we examined the expression of *Hes1* in control and *Piezo^dbKO^* mice. ScRNAseq analysis showed an overall 1.4-fold increase of *Hes1* and a 2-fold increase specifically in the absorptive progenitors, cluster 0 (**Fig. 4A**). Interestingly, scRNAseq and smFISH labeling both exhibited a compelling decrease of *Atoh1*-expression in the absence of PIEZO (**Fig. 4B**). These results suggest that PIEZO channels are essential to temper NOTCH signaling in maintaining secretory cell lineage specification. To further analyze NOTCH activity, we treated *Hes1-GFP* organoids with GsMTx4 and found a strong increase in the GFP signal, confirming that PIEZO inhibits NOTCH activation (**Fig. 4C**).

**FIGURE 4:**
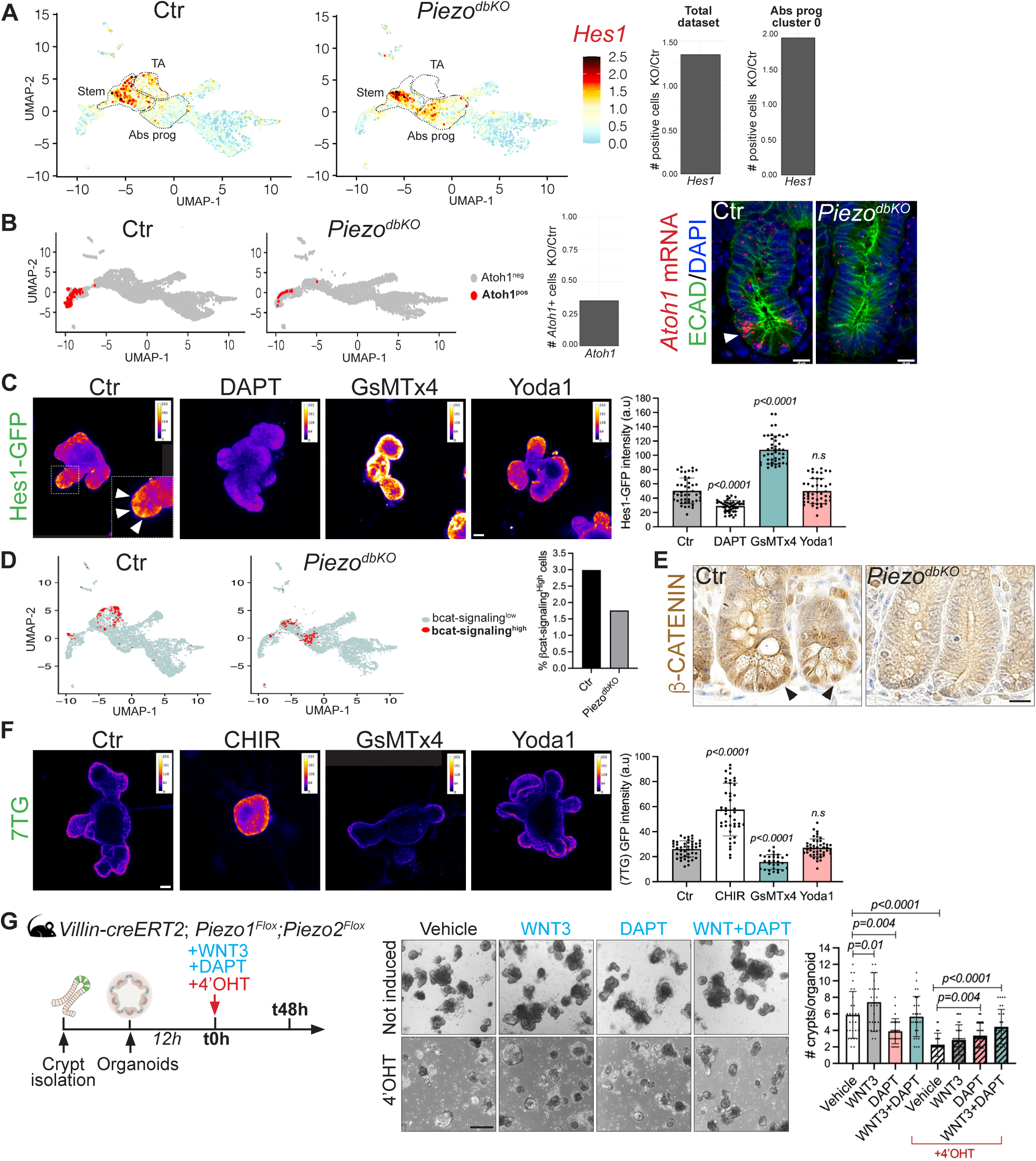
PIEZO mechanotransduction regulates the NOTCH and WNT pathways. **(A)** UMAP plot of *Hes1* expression Ctr and *Piezo^dbKO^* mice. Fold change expression of *Hes1* from *Piezo^dbKO^* over Ctr mice in all intestinal epithelial cells (left graph) and in absorptive progenitor cluster 0 (right graph). **(B)** UMAP plot of *Atoh1* in Ctr and *Piezo^dbKO^* and fold change expression over Ctr. Representative images of smFISH for *Atoh1* in Ctr and *Piezo^dbKO^* crypts. ECAD immunostaining (green) shows epithelial cells. **(C)** Representative images of organoids expressing the NOTCH reporter Hes1-GFP exposed to DAPT (NOTCH inhibitor), GsMTx4 and Yoda1. Pseudo-color shows intensity of the GFP reporter fluorescence quantified on the right panel. Arrows in the inset indicate GFP^high^ cells in the crypt (n=45 crypts/condition; N=3 independent experiments). **(D)** UMAP plot of cells expressing βcatenin-signaling gene signature (threshold >0.1= bcat-signaling^High^ cells) in Ctr and *Piezo^dbKO^* crypts. Quantification of the percentage of bcat-signaling^High^ cells in Ctr and *in Piezo*-KO mice. **(E)** β−CATENIN immunocytochemistry in Ctr and *Piezo*-deleted intestines 4 days post-tamoxifen induction. Arrowheads showing nuclear β−CATENIN signal. Scale bar, 25μm. **(F)** Representative images of organoids expressing the WNT reporter 7TG exposed to CHIR (WNT activator), GsMTx4 and Yoda1. Pseudo-color shows intensity of the GFP reporter fluorescence quantified on the right panel (n=45 crypts/condition; N=3 independent experiments). **(G)** Experimental scheme of *Piezo^dbKO^* organoids induced *in vitro* with 4’OHT and treated with WNT3, DAPT or both. Brightfield representative images of WNT3, and DAPT-treated *Piezo^dbKO^* organoids with and without 4’OHT induction. Quantification of organoid growth as the number of crypt/organoid (n=30 organoids/condition; N=3 independent experiments). Scale bar, 10μm (B and D) and 25μm (C and E). Error bars indicate standard deviation (SD). Mann-Whitney test (C, F), Two-tailed paired Student’s *t*-test (G). Box edges show 25th and 75th percentiles, central point is median, and error bars represent minimum and maximum values. *P* values are shown in panels. See also Supplementary Figure S5 and S6.

We then analyzed WNT signaling, the main driver of stem cell proliferation and maintenance. Notably, within the niche, the threshold of WNT signaling is critical, and the pathway engages several feedback loops that balance the opposing processes of cell differentiation and self-renewal (*30*). ScRNAseq data analysis revealed that the percentage of cells expressing βcatenin-signaling gene signature (*31*) was decreased in *Piezo*-deleted crypt cells (**Fig. 4D**) which is consistent with the absence of nuclear β-CATENIN in *Piezo^dbKO^* mice (**Fig. 4E**). To monitor WNT activity, we generated the WNT signaling reporter 7TG (7x-Tcf-eGFP) organoid line using lentiviral transduction **(Fig. S6A**). 7TG organoids treated with the PIEZO inhibitor GsMTx4 depicted a significant decrease of GFP signals (**Fig. 4F**). Surprisingly, overactivation of PIEZO with Yoda1 did not affect NOTCH or WNT signaling activity (**Fig. 4C, 4F**), nor did it impact organoid growth and *de novo* crypt formation (**Fig. S6B, Movie S4**), suggesting that basal PIEZO activity is sufficient to regulate downstream pathways and cell fate choice. We then hypothesized that restoring levels of NOTCH and WNT pathways could rescue ISC function in the absence of PIEZO. We therefore ablated *Piezo* expression in organoids and cultured them in presence of a NOTCH inhibitor, DAPT, and/or WNT3 ligand (**Fig. 4G**). Intriguingly, the combination of NOTCH inhibition and WNT activation restored organoid growth (**Fig. 4G**).

Taken together, these data indicate that PIEZO channels are required to coordinate stem cell proliferation, self-renewal and differentiation of downstream NOTCH and WNT pathways.

### Different mechanical stimuli in the intestinal crypt activate PIEZO channels *ex vivo*

In addition to these downstream signaling mechanisms, we sought to identify the upstream stimuli that activate PIEZO channels in stem cells. PIEZO1 has been shown to sense the local substrate stiffness to modulate human neural cell fate choice (*13*). Moreover, *in vitro* experiments demonstrated that the size of the stem cell compartment and the number of ISCs decrease with substrate rigidity (*23*). To test whether ISCs sense the stiffness of their niche through PIEZO channels, we cultured organoids as 2D monolayers seeded on PAA gels of different stiffnesses. We first evaluated whether the 2D monolayer was a suitable model for our study by generating monolayers from 3D crypt organoids of non-induced *Piezo^dbKO^*mice (**Fig. S7A**). Transcript analysis of *Piezo1* and *Piezo2* 48h after *in vitro* induction with 4’OHT established successful recombination efficiency in *Piezo^dbKO^* monolayers (>95%; **Fig. S7B**). Importantly, 4’OHT-treated monolayers showed a 90% decrease in ISCs (**Fig. S7C**) and 80% in Paneth cells (**Fig. S7D**). Although both proliferation and absorptive cells (EZRIN+) were unaffected by *Piezo* loss (**Fig. S7E, S7G**), mutant monolayers showed a decrease in secretory cell numbers (UEA+ cells (**Fig. S7F**). These results recapitulated the *in vivo* mouse phenotypes described at day 6 post-tamoxifen treatment (**Fig. 2**) and validated the 2D monolayer as a reproducible and suitable *ex vivo* model for studying the role of PIEZO channels in ISCs.

Several studies have shown that PIEZO function increases intracellular Ca^2+^ in response to stiffness (*13*, *32*). To assess how PIEZO activity is modulated by stiffness in ISCs, we generated a new Lgr5-GFP+ organoid line with stable overexpression of the FusionRed-based Ca^2+^ reporter K-GECO1 to visualize Ca^2+^ influx in real-time using live imaging (*33*). Because the calcium baseline in Lgr5-GFP+ crypts was low (**Fig. S7H**), we induced PIEZO activation using Yoda1 to facilitate the analyses of calcium transients (*34*). We found that PIEZO-mediated increase in intracellular Ca^2+^ levels in ISCs scaled with stiffness, with higher transients in monolayers cultured at 18kPa compared to 1.5kPa (**Fig. 5A-C** and **Movies S5-S6**). This suggests that PIEZO is more prone to activation on a stiff substrate; consequently, its contribution to the regulation of ISC maintenance may depend on the niche stiffness. To test this hypothesis, we cultured monolayers expressing membrane td-Tomato on gels of increasing stiffnesses: 1.5kPa, 5kPa, 11kPa, 18kPa and 30kPa (**Fig. 5D**). To investigate whether PIEZO channels are involved in ISC stiffness sensing, we either blocked (GsMTx4) or activated (Yoda1) PIEZO channels using pharmacological drugs (**Fig. 5D**). Consistent with a recent report (*23*), the stem cell compartment was reduced with increasing stiffness (**Fig. 5E, F** and **Fig. S7I**). Interestingly, on soft substrates (1.5kPa to 5kPa), PIEZO activation with Yoda1 enhanced ISC area. At the same time, its inhibition had no effect (**Fig. 5E, F** and **Fig. S7I)**, suggesting that the PIEZO activity was already minimal on low rigidity and cannot be further inhibited. On the contrary, activation of PIEZO on stiffer substrates (11kPa to 30kPa) showed no difference in ISC area, while its inhibition significantly diminished the stem cell compartment (**Fig. 5E, F** and **Fig. S7I)**. These results suggest that ISCs at the bottom of the crypt could experience a stiffer environment that allows PIEZO activation. To test this *in vivo*, we measured the stiffness around the basement membrane at the bottom and top of the crypt using Atomic Force Microscopy (AFM) on snap-frozen intestinal tissue slices (**Fig. 5G-H**). We found that the average elasticity at the basement membrane region of the crypt bottom was higher than the elasticity at the top (**Fig. 5I**), revealing that ISCs reside in a stiffer environment than the cells at the top of the crypt. Altogether, these results reveal that PIEZO channels are more prone to activation by a stiff substrate, such as the crypt bottom, and are required for a proper response to stiffness-dependent changes in ISC function.

**FIGURE 5:**
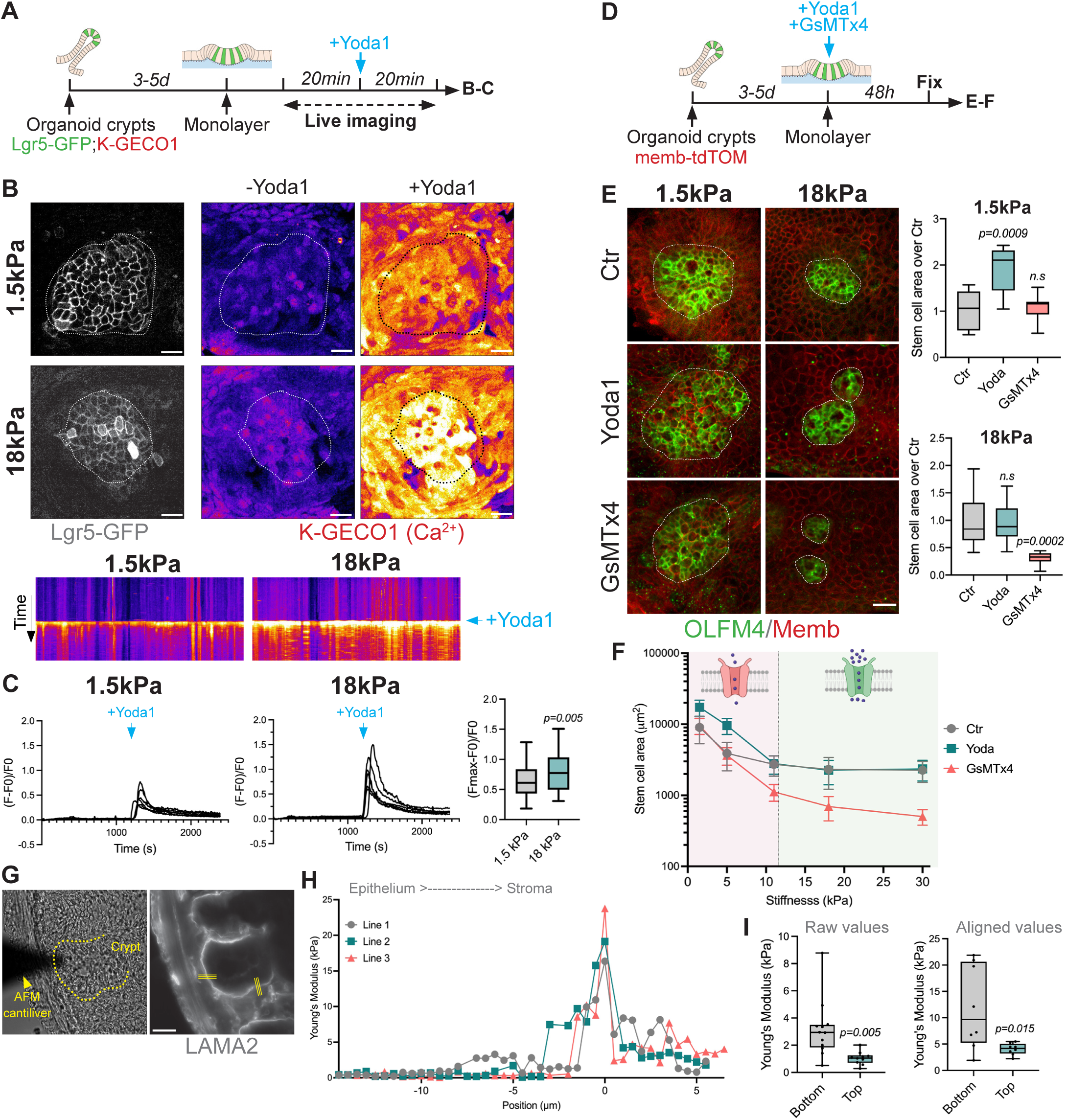
Modulation of niche stiffness activates PIEZO channels to promote stemness. **(A)** Experimental scheme of Ca^2+^ live imaging as a proxy for PIEZO activity using Lgr5-GFP;K-GECO1 organoid monolayers plated on 1.5kPa (soft) and 18kPa (stiff) PAA gels. To visualize efficient Ca^2+^ transients, monolayers were treated with Yoda1 PIEZO activator. Monolayers were imaged for 20min (5sec intervals) before and immediately after adding Yoda1. **(B)** Time series of calcium influx (FIRE LUT) in Lgr5-GFP+ crypts in organoid monolayers plated on 1.5kPa and 18kPa gels before (-) and after addition (+) of the PIEZO activator, Yoda1. Kymographs from the time-lapse Movies S4 (1.5kPa) and S5 (18kPa) before and after Yoda1 induction. X axis represents individual cells. Dashed line delineates crypt region. Scale bar, 25μm. **(C**) Calcium (K-GECO1) traces and max intensity increase of calcium (dF_max_/F_0_) after Yoda1 in Lgr5-GFP+ crypts (n=45 crypts from N=3 independent experiments). Each trace represents an individual Lgr5-GFP+ crypt measurement and n=6 crypts/condition are depicted per graph. **(D)** Experimental scheme of PIEZO channel pharmacological inhibition (GsMTx4) or activation (Yoda1) on 2D organoid monolayers. Monolayers were formed 3-5 days after seeding organoid crypts expressing the membrane tdTomato on substrates of different stiffnesses (1.5kPa, 18kPa). Stem cell area was analyzed 48h after PIEZO inhibition/activation. **(E)** Representative images of OLFM4 immunostaining and the membrane tdTomato (Memb) of monolayers cultured on substrates of 1.5kPa and 18 kPa and treated with PIEZO inhibitor (GsMTx4) or activator (Yoda1). Quantification of the stem cell area over non-treated control (Ctr) (n= 15 crypts from N=3 independent experiments). Dashed line delineates crypt region. Scale bar, 25μm. **(F)** Stem cell area of monolayers cultured on substrates of increasing stiffnesses showing that PIEZO channels are more prone to activation on stiffer substrate (>11kPa; “PIEZO ON”) and less active on soft substrate (<1.5kPa; “PIEZO OFF”). **(G)** Image of AFM measurement showing the cantilever and the intestinal crypt. LAMININ alpha 2 (LAMA2) immunostaining labelling the crypt basement membrane and the probed regions for AFM measurements (yellow lines). Scale bar, 25μm. **(H)** Aligned Young’s Modulus traces for the 3 lines represented in panel (G). **(I)** (left) Quantification of the Young’s Modulus (average of all values) for all bottom and top crypt datasets. (right) Quantification of the Young’s Modulus (average of peak values) for the selected bottom and top crypt datasets. Error bars indicate standard deviation (SD). Mann-Whitney (C), Kolmogorov-Smirnov test (E), Two-tailed unpaired Student’s *t*-test (I). Box edges show 25th and 75th percentile, central point is median, and error bars represent minimum and maximum values. *P* values shown in panels; *n.s*: non-significant. See also Supplementary Figure S7.

Another potential PIEZO activator is tissue tension (*11*). Previously, we demonstrated that the bottom of the villus is under tension that dissipates towards the villus tip (*35*). Moreover, we also showed that tension within the crypt builds up towards the stem zone (center of the crypt) and decreases at the TA zone (*23*). Thus, we hypothesized that ISCs sense the changes in tension through PIEZO, which induces cell fate decisions. To test this, we engineered a cell stretching device to modulate tension in monolayers by applying uniaxial stretch using a traction motor (**Fig. 6A**). Given that PIEZO channels were less prone to activation on soft substrates (**Fig. 5D**), we plated monolayers on 5kPa gels and applied a continuous cyclic stretch (30sec stretch, 30sec release, for 24h, 10% strain, 0.26Hz) (**Fig. 6B**). Interestingly, inducing exogenous stretch to monolayers increased the OLFM4+ stem cell area by 3-fold (**Fig. 6C**), suggesting that mechanical stimulation regulates ISC behavior. To investigate whether PIEZO drives this phenotype, we blocked PIEZO activation using GsMTx4 while stretching (**Fig. 6B**). Inhibition of PIEZO decreased the OLFM4+ stem cell area but not to the extent of the non-stretched control level (**Fig. 6C**), implying that an additional, PIEZO-independent mechanism, might be involved in ISC mechano-sensing of the niche. Interestingly, stretching induced no change in the apical area (**Fig. 6D**) but increased the number of EdU^+^ cells (**Fig. 6E**), suggesting that stem cell amplification is not due to increase of cell size but through proliferation. These data demonstrate that ISCs transduce mechanical changes of the microenvironment through PIEZO mechanosensitive channels to regulate their cell fate decision according to the biomechanical properties of the stem cell niche.

**FIGURE 6:**
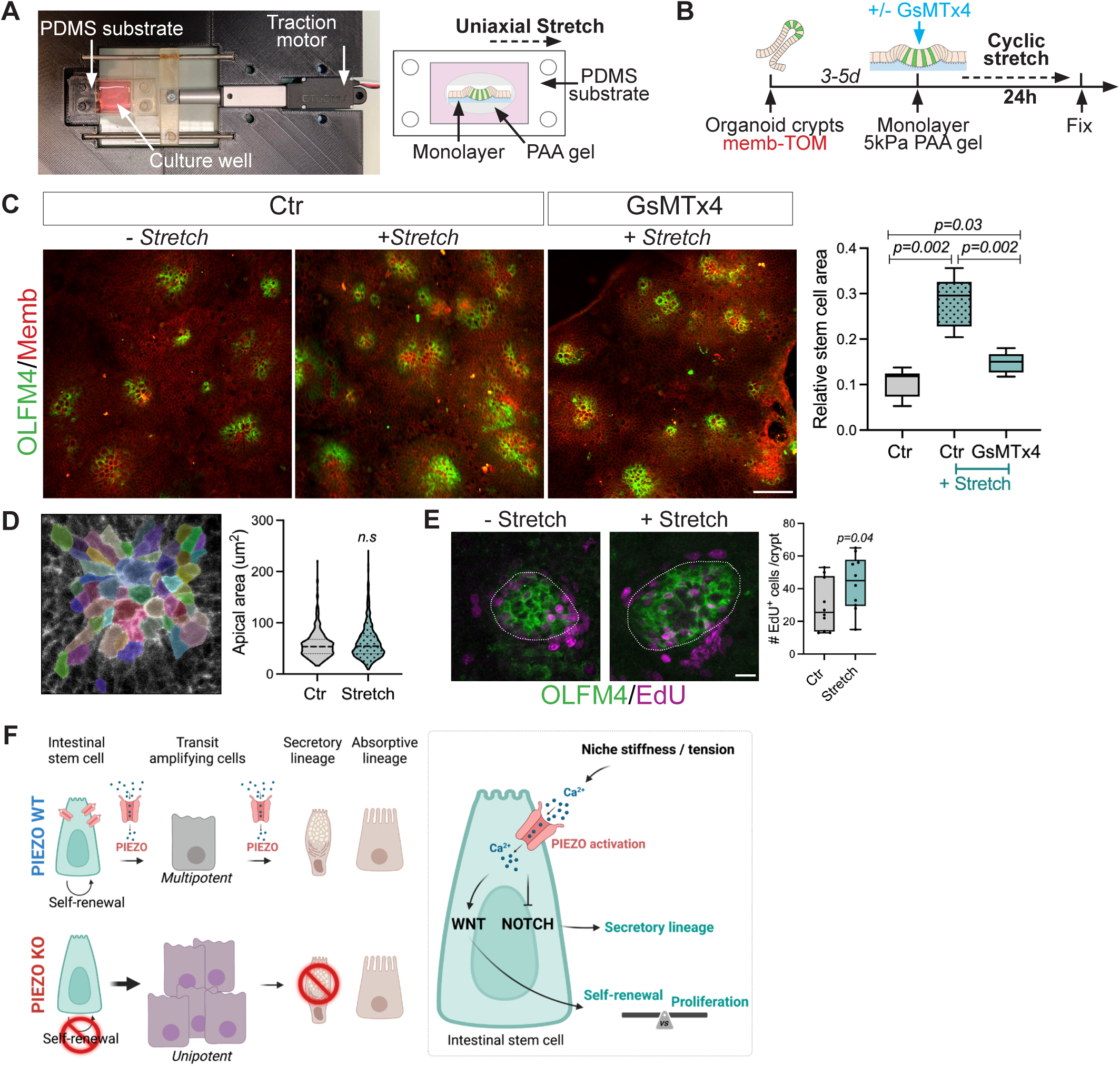
PIEZO channels respond to tissue tension to modulate ISC function. **(A)** Image and schematic of the cell stretching device. 2D monolayer was seeded on a polyacrylamide (PAA) gel attached to a PDMS substrate that undergoes a given uniaxial stretch using a traction motor. **(B)** Experimental scheme of monolayer seeded on a 5kPa PAA gel and stretched for 30sec for 24h, 10% strain, 0.26Hz. 2D monolayer were either non-treated (Ctr) or treated with PIEZO inhibitor GsMTx4. **(C)** Representative images of OLFM4 immunostaining and membrane tdTomato (Memb) of control (Ctr) and GsMTx4-treated monolayers with (+) or without (-) cyclic stretch (n= 45 crypts from N=3 independent experiments). Scale bar, 50μm. **(D)** Representative segmentation mask (Cellpose v.2 (*64*)) of crypt cells on the z plane that corresponds to the apical side of cells using membrane-TOM staining. Quantification of apical area with or without stretch (n= 1531 cells, from n=35 crypts, 3 monolayers). **(E)** OLFM4 immunostaining and EdU labelling (2h) in crypts from monolayer with or without stretch. Quantification of the number of EdU+ cells/crypt (n=20 crypts, N=3 independent experiments). Dashed line delineates crypt region. Scale bar, 10μm. **(F)** Proposed model for PIEZO mechano-sensing of the intestinal stem cell niche. In intestinal stem cells, PIEZO channels are required to transduce mechanical stimuli from the extracellular niche to intracellular signal, leading to cell fate decisions. Conditional KO of PIEZO in intestinal epithelial cells induces self-renewal and secretory lineage specification defects, leading to ISC and secretory cell loss. Mechanistically, upon PIEZO channel activation by increase in niche stiffness and/or tension, intracellular Ca^2+^ influx is generated, leading to the repression of NOTCH signaling for secretory cell differentiation and the modulation of WNT signaling for the maintenance of proper self-renewal vs proliferation balance. Error bars indicate standard deviation (SD). Kolmogorov-Smirnov test (C), Mann-Whitney (D, E). Box edges show 25th and 75th percentile, central point is median, and error bars represent minimum and maximum values. *P* values shown in panels.

## DISCUSSION

Unraveling the mechanisms by which stem cells integrate mechanical cues from their microenvironment is essential for understanding tissue homeostasis. Here, we propose a comprehensive model for PIEZO mechano-sensing of the intestinal stem cell niche (**Fig. 6F**). In ISCs, PIEZO channels are required to transduce mechanical stimuli from the extracellular niche into intracellular signals, regulating stem cell fate decision and maintenance. Conditional KO of PIEZO in intestinal epithelial cells induces self-renewal and lineage specification defects, leading to ISC and secretory cell loss. Mechanistically, upon PIEZO channel activation by niche stiffness and/or tension, intracellular Ca^2+^ influx is generated; subsequently, this activation represses the NOTCH pathway to induce secretory cell differentiation and modulates WNT signaling to maintain appropriate self-renewal *vs.* proliferation balance (**Fig. 6F**). Furthermore, our rescue experiments (**Fig. 4G**) demonstrated that WNT and NOTCH signaling act synergically to maintain ISC function downstream of PIEZO mechanotransduction activity. Interestingly, these findings align with those of a previous report demonstrating that NOTCH antagonizes WNT signaling, maintaining it at levels necessary for the proper simultaneous maintenance of stem cells, proliferation and differentiation (*36*).

It is well established that PIEZO mechanotransduction mechanism permeates calcium that further modulates downstream intracellular pathways (*37*). Indeed, using a calcium reporter organoid line, we showed that PIEZO activation induces intracellular Ca^2+^ influx both in 2D monolayers (**Fig. 5B**) and in 3D organoids (**Fig. S8 and Movies S7, S8**). Interestingly, it has been demonstrated that increase of cytosolic Ca^2+^ in both mouse and human cell lines (*38*, *39*), and in the fly midgut (*14*) inhibits NOTCH activity further confirming the direct link between PIEZO-induced calcium and NOTCH pathway. At the mechanistic level, it has been shown that calcium regulates NOTCH by stabilizing he Notch-extracellular domain with the transmembrane domain, and that disruption of Ca^2+^ (with EDTA) leads to shedding of the NOTCH intracellular domain (NICD), which translocates to the nucleus to activate the transcription of target genes (*40*). Consequently, we speculate that PIEZO-induced Ca^2+^ in ISCs tempers NOTCH by stabilizing the receptor heterodimer, thereby preventing its cleavage and subsequent pathway activation. Regarding WNT, it has been shown that hydrostatic pressure activates PIEZO in odontoblasts to promote their differentiation through activation of the WNT pathway (*41*). Also, WNT/β-catenin, known to be essential for osteoblast differentiation, is diminished in *Piezo* KO osteblasts during bone development (*42*). But, how Ca^2+^ modulates WNT signaling at the biochemical level is still unclear.

We further described the upstream mechanical cues that induce PIEZO activation. We showed that PIEZO channels were more prone to activation on rigid substrates with stiffness values ranging from 11kPa to 30kPa (**Fig. 5E-F** and **Fig. SS7I**). Although our AFM measurements indicated that the stiffness of the basement membrane region at the bottom of the crypt is of the same order of magnitude (between 2-20kPa), the strict comparison could not be made due to the limitation of AFM measurements. Nevertheless, these *in vivo* AFM measurements showed that ISCs reside in a stiffer environment at the bottom of the crypt (**Fig. 5I**) which is consistent with our experiments on 2D monolayers. Moreover, the 2D organoid monolayers cultured on PAA gels with different stiffnesses also provide essential mechanistic knowledge on how ISCs respond to alterations in their microenvironment rigidity. Consistent with our data, a recent study using 2D intestinal organoids grown on a hydrogel matrix showed that increased stiffness decreased the Lgr5+ ISC number (*23*, *43*). Interestingly, on a soft substrate (1.5-5kPa), PIEZO inhibition does not affect the ISC area, suggesting that at low rigidity, PIEZO is not involved in their regulation.

Regarding tension, we previously showed that villus tension is generated by active cell migration being highest at the villus base and decreasing towards the tip of villi (*44*). Within the crypt, tension builds up from the crypt edge towards center, where stem cells are located (*23*). However, the exact origin of this tension within the crypt remains unknown. We speculate that the stretching of cells could be attributed to several factors: the division of neighboring epithelial cells, contractile fibroblasts located just beneath the basement membrane and/or peristalsis - the contraction of the muscle that in proximity to the stem cell niche. For that reason, we intended to mimic tension induced by peristalsis, described as approximately 16 contractions per minute (0.26Hz), using our stretching device on 2D monolayers (**Fig. 6**). Stretching induces an increase in stem cell area (**Fig. 6C**) without changes in cell size (**Fig. 6D**). These results suggest that peristalsis could be one of the mechanisms for promoting stemness PIEZO channels activation. Corroborating the previous report (*17*), we demonstrated that the ISC compartment expanded with mechanical stretch through proliferation and this expansion was partially blocked by PIEZO inhibition. These results suggest that ISCs might employ another PIEZO-independent mechanism, such as integrin-mediated mechanotransduction (*45*), to transduce niche biomechanical cues. We found that PIEZO1 was predominantly located in crypt cells (**Fig. 1C**). Thus, we mainly focused on the effect of stretching on crypt cells and specifically ISCs; however, it is likely that other epithelial cells also respond to changes in tension and adapt their behavior accordingly. Interestingly, it has recently shown in 3D organoids that PIEZO activity was necessary for inflation-mediated fission by activating stretch-dependent downstream calcium signaling. This study further confirms the involvement of PIEZO channels in stretch-related mechanisms in intestinal epithelial cells (*46*).

Recent studies have demonstrated that crypt topology and cell geometry drive epithelial cell patterning (*47*) and regulate ISC self-renewal and function (*48*). In the 2D monolayer, the curvature of the crypt domains is diminished with the substrate rigidity (*23*). However, the curvature is similar between 5kPa and 11kPa (*27*), while the PIEZO response differed. Thus, it is unlikely that crypt topology triggers PIEZO activation in ISCs. Another possibility could be that the PIEZO channel senses stimuli other than rigidity and membrane tension. It has been shown that PIEZO1 channels in the *Zebrafish* intestine are important sensors of cell crowding and control the number of cells in the epithelia by regulating cell extrusion events (*49*). A recent study reported that cell density and lateral crowding play crucial roles in stem cell fate decisions in the embryonic epidermis (*50*). Thus, cell crowding within the intestinal crypts, as a consequence of proliferation, could activate PIEZO channels in ISCs, and this remains to be addressed. Similarly, volumetric compression increases intracellular crowding and promotes organoid growth by enhancing WNT signaling (*51*).

Finally, given the critical requirement of PIEZO channels in ISC maintenance, it would be of great interest to understand their potential roles in aging, upon repair, or in pathological contexts such as chronic inflammation.

## Supporting information

supplementary txt

## MATERIALS AND METHODS

### Mice used in this study

Animal housing, husbandry and handling were approved and performed in accordance with the Animals for Research Act of Ontario and the Guidelines of the Canadian Council on Animal Care at The Toronto Centre for Phenogenomics. The health and immune status of all mice used were normal. Food and water were administered *ad libitum*. Comparisons were made between age-matched littermates using 8-12 weeks-old mice. As no phenotype difference was observed between males and females, mice of both sexes were used in all experiments. *Villin-*creERT2 (*52*) and R26^mTmG^ (*53*) were purchased from The Jackson Laboratory while mice. *Piezo1^flox^*, *Piezo2^flox^* were a kind gift from Dr. Huang (SickKids). *Hes1-GFP;R26^mTom^* used to generate crypt organoids were a gift from Dr. Fre (Institut Curie).

### Tamoxifen and EdU administration

Mice were injected intraperitoneally once a day with 100μl of tamoxifen (20mg/ml) for indicated times. To assess *in vivo* proliferation, mice were injected with 5-ethynyl- 2′-deoxyuridine (EdU, 30μg/g mouse, Sigma 900584) 2h before sacrifice. *In vitro,* recombination was induced by treatment with 10μM 4-hydroxytamoxifen (4’OHT; H7904) in methanol for 48h, and the efficiency of recombination was determined by RT-qPCR.

### Histology, Immunostaining, in situ hybridization (smFISH) and TUNEL staining

Whole intestines and colons were dissected, flushed gently with cold 1X PBS to remove fecal content, and fixed overnight at 4°C in 4% paraformaldehyde in 1X PBS. After three washes in 1X PBS for one hour, tissues were dehydrated in ethanol and embedded in paraffine. Sections (7μm) were deparaffinized and rehydrated using standard techniques (xylene and descending ethanol gradients). Histology was assessed by staining with Harris hematoxylin/Eosin Y or Alcian Blue/Nuclear Red.

For immunostaining, antigen retrieval was performed by incubating sections in boiling 10mM sodium citrate pH 6 for 20min in a steamer. After three washes in 1X PBS for one hour, the sections were permeabilized and blocked in a buffer containing 0.25% Triton X-100 (Sigma) and 10% goat serum (Gibco) for 1h at RT. The tissues were then incubated with primary antibodies overnight at 4°C. The following primary antibodies were used in this study: rabbit anti-LYSOZYME (1:1000, Dako, A0099), rabbit anti-OLFM4 (1:200, Cell Signalling, 39141S), mouse anti-ECADHERIN (1:300, BD, 610182), EZRIN (home-made), DCLK1 (1:200, Cell signaling, 62257S), MUC2 (1:200, Genetex, GTX100664), Phospho-histone 3 (1:200, sigma, H0412), ANPEP (1:300, R&D System, AF2335-SP), ChgA (1:200, Abcam, ab85554). The sections were washed with 1X PBS three times and incubated with Alexa-conjugated secondary antibodies (Life Technologies, 1:500) and Hoechst (Life Technologies, 1:10000) for 45 min at RT. EdU staining was performed using the Click-iT Plus kit (Life Technologies, C10640) according to the manufacturer’s recommendations

For PIEZO immunostaining, slides were incubated in ice-cold MetOH at −20°C for 10min prior to permeabilization and blocking as described above. PIEZO antibody (Novus, NBP −78446SS, lot D147738, 1:100) was diluted in the blocking buffer and incubated at RT for 2h.

β−CATENIN (1:50, BD, 610154) immunohistochemistry was performed as described before (*54*). Briefly, 7μm sections were incubated in boiling 40mM Tris 1mM EDTA pH9 antigen retrieval solution for 20min. After blocking in 1% BSA/0.5% triton for 30min at RT, primary antibody was incubated 2h at RT in 0.05% BSA. Secondary biotinylated antibody (1:150, Novus) was incubated 1h at RT followed by ABC HRP kit (Vector labs, PK6100) and DAB Substrate kit (Sigma, 34002) according to manufacturers’ recommendation.

Single molecule FISH (smFISH; ACD, RNAscope® Multiplex Fluorescence Detection Kit v2 (323110) was performed according to manufacturer guidelines using *Lgr5* (312171), *Atoh1* (408798), *Axin2* (400331), *Tff3* (573078), *Muc2* (318831), *Fabp1* (562831), *Alpi* (515391), *Piezo1* (500511) and *Piezo2* (400191) probes. Each experiment includes a positive (320881) and negative (320871) control probe to validate signals.

TUNEL staining has been performed using the ApoTag^®^ Peroxidase *In Situ* Apoptosis detection kit (Sigma, S7100) following the manufacturer’s guidelines.

Histology images were acquired using Nikon microscope with DS-Fi2 camera using a 10X/0.3 objective. Confocal images were acquired with Zeiss LSM880 microscope using 25X/0.8 objective and Zen software. Quantifications were done using Fiji cell counter plugin.

### Intestinal 3D organoids

Organoids from crypts were performed as described previously (*55*). Briefly, isolated crypts were plated in 50% growth-factor free Matrigel (Corning, 354234) and cultured with ENR media (Advanced DMEM/F12 (Gibco)/ 1X Glutamax (Gibco)/ 1% penicillin and streptomycin (PS; Gibco)/ 10 mM HEPES (Sigma, H0887)/ 1X B-27 (Gibco)/ 1X N-2 (Gibco), 50 ng/ml of human EGF (Sigma, E9644)/ 100 ng/ml Noggin (Cerderlane labs, 6057NG)/ 500 ng/ml of mouse R-spondin-1 (RnD, 3474RS)/ 1 μM N-acetyl-l-cysteine (Sigma-Aldrich). Organoid forming capacity (= number of organoids formed/number of plated crypts) was determined after two days of culture, and the number of crypts per organoid was assessed by the number of buds per organoid. Primary organoids were cultured for 7 days before passaging. Subculturing secondary organoids was performed by mechanically disrupting primary organoids with a 18G needle until full dissociation. Live imaging was performed using Zeiss Cell Discoverer 7 using 5x/0,35 objective for 36h with one image per hour.

Organoids were treated with 20 μM gadolinium chloride hexahydrate (GdCl3. 6H2O) (Sigma, G7532), 10 μM GsMTx4 (Biotechne, 4912), 30 μM Yoda1 (Biotechne, 5586), 3 μM CHIR-99021 (Euromedex, AB-M1989), 500nM DAPT (Tocris, 2634), 200 ng/ml WNT3 (Peprotech, 315-20) as indicated in figure legends.

For immunostaining, 3D organoids were fixed with 4% PFA 1h at RT. After two PBS1X washes, they were treated with 1% Triton/10% goat serum for 1h at RT, washed twice and incubated with primary antibody in PBS/0.05% triton/10% goat serum overnight at RT. After 2 washes of 30min at RT, secondary antibody was incubated 5h at RT.

For smFISH, organoids were fixed and permeabilized in 1% Triton as described above, then treated with H_2_0_2_ 15min and Protease Plus reagent for 30min at 40°C. The rest of the protocol was performed according to manufacturer’s protocol.

### Flow cytometry and analysis

Organoids flow cytometry gating and analysis have been performed as described here (*55*). Briefly, organoid cultures were recovered using ice-cold Cell Recovery™ Solution (Corning, C354253) and incubated in TrypLE Express (Gibco) with 1,000 U/ml of DNaseI (Roche, 11284932001) for 3 min at 37°C. Cells were washed with (Hank’s Balanced Salt Solution (HBSS, Gibco)/ 2% fetal calf serum (FCS; Gibco)/10mM HEPES and stained with the following antibodies: CD24-Pacific Blue (Biolegend, 101829, 1:500) and Epcam-PE-Cy7 (Biolegend, 324221, 1:500) and cells were supplemented with propidium iodide (Invitrogen, P1304MP) to exclude dead cells. Data were recorded using BD LSRFortessa and FACSDIVA software. Analysis was performed with FlowJo software.

### RNA extraction and quantitative PCR (RT-qPCR)

RNA extraction from organoids has been thoroughly described here (*55*). Briefly, organoids were dissociated from Matrigel using ice-cold Cell Recovery™ Solution (Corning, C354253), recovered by centrifugation 160g for 10 min, resuspended in Qiagen RLT^®^ buffer/1% b- mercaptoethanol and RNAs were extracted using Qiagen RNeasy^®^ Plus Mini extraction kit according to the manufacturer’s instructions. Two wells were combined per data point. Reverse transcription was performed using SuperScriptIII (Thermo, 18080093) according to the manufacturer’s protocol. The expression of mature mRNAs was assessed with Power SYBR green master mix (Roche, 04913914001) and run on QuantStudio 5 (Applied Biosystems). The analysis was performed using the 2^-ΔΔCT^ method (*56*). All RT-qPCR samples were normalized with *Tbp* and *Rpl13* expression values. Specific forward and reverse primers used for RT-qPCR are listed in supplementary Table S1.

### Generation and analysis of 2D organoid monolayers

Monolayers were generated as described here (*23*). Glass-bottom dishes (Fluorodish) were incubated for 25min at RT with a solution of Bind-Silane/absolute EtOH/acetic acid at a volume proportion of 1:12:1. After two washes with absolute EtOH, 12μl of the polyacrylamide (PAA) mix were added on the glass and covered with a 12mm round coverslip (stiffnesses recipes are in Supplementary Table 2). After 1h of polymerization at RT, the coverslip was removed, and gels were washed with PBS1X. For surface activation, gels were treated with 2 mg/ml Sulpho-SANPAH (Sigma, 803332), irradiated for 7.5 min with ultraviolet light (365 nm) and washed twice with 10 mM HEPES (Gibco) under agitation. Gels were then coated with 250μg/ml Collagen I (First Link UK) and 100μg/ml Laminin I (Thermo, 23017015) overnight at 4°C.

3D organoids were recovered using an ice-cold Cell Recovery™ Solution and mechanically broken by pipetting 15 times with a syringe equipped with a 18G blunt needle. Organoids were centrifuged at 160g, 10min at 4°C and the pellet was resuspended in ENR supplemented with 10μM Y-27632 (ATCC, ACS-3030). Four organoid-containing wells of a 24-well plate were pooled in 50μl of ENR, seeded on one ECM-coated PAA gel and incubated at 37°C, 5% CO_2_. After 4h, 550μl of pre-warmed ENR/Y-27632 medium was added to the dish and monolayers were analyzed 3-5 days after seeding. Of note, Y-27632 is removed from the medium after 3 days of culture.

For generation of 2D monolayer from single cell suspension, 3D organoids were dissociated as described before (*57*). Briefly, organoids were harvested from Matrigel as described above. For cell dissociation, the pellet was suspended in AccuMax (Sigma, A7089) containing CHIR-99021, Y-27632 (CY), incubated at 37°C for 8 min and then digestion was stopped adding by DMEM- F12 containing B27 and CY. Organoids were further mechanically dissociated by pipetting up and down and then centrifuged at 400 × *g* for 5 min at 4°C. Single cell suspension was resuspended in 50µl of ENR-CY per PAA gel by mixing control and mutant organoids at 1:1 ratio. ENR-CY was replaced by ENR 2 days later.

For RNA extraction, monolayers were scrapped out of the PAA using a cell scraper and 350μl Qiagen RLT^®^ buffer/1% þ-mercaptoethanol and RNAs were extracted using Qiagen RNeasy^®^ Plus Micro extraction kit according to the manufacturers’ instructions. RT-qPCR was performed as described above.

For immunostaining, 2D organoid monolayers were fixed in 4% paraformaldehyde (Electron Microscopy Sciences) for 10min at RT, washed 3 times with PBS1X and blocked with a solution containing 0.25% Triton X-100 (Sigma)/10% Goat (Gibco)/PBS1X for 1h. Primary antibodies diluted in the blocking solution were added and incubated overnight at 4°C in a humid chamber. After 3 washes with PBS1X, secondary antibodies (1:500) were added for 45min at RT. Ulex Europaeus Agglutinin I (UEA I), DyLight® 649 (1:1000, Sigma, DL-1068) was incubating along with the secondary antibody.

### Stretching device

The stretcher system comprises a linear servo (L12-30-210-6-R, Actuonix) connected to an ATmega328 microcontroller (Arduino Nano). The servo is directly powered by the Arduino’s 5V pin, the commands are sent from a computer using the Arduino’s serial interface by an open-source protocol (https://github.com/araffin/arduino-robust-serial/tree/master). The PDMS chips are attached using two 3D printed parts (resist: RG35B, BASF; printer: 028J+ HR, DWS): one at head of the servo and the other on the breadboard (MSB30/M, Thorlabs) (*58*). PDMS substrates were fabricated using PDMS 10:1 (monomer/crosslinking agent) poured in a metal mold and incubated at 90°C for 2h. The resulting stretchable PDMS substrates were peeled off the mold and UV irradiated in a plasma cleaner for 1 min as described here (*59*). To activate the surface of the PDMS well, we added 10% APTES (3-Aminopropyl triethoxysilane; Sigma) in absolute ethanol for 1 h at 65 °C. After extensive cleaning with PBS1X, the well was incubated with 1.5% Glutaraldehyde/PBS1X for 25 min at room temperature. After cleaning with PBS1X, treated PDMS substrates were air-dried before use. PAA hydrogels were polymerized between two coverslips treated with Repel-Silane (2% dimethyldichlorosilane). After 1h of polymerization, one coverslip was removed and the PAA gel was pressed against the coated stretchable PDMS well for 10sec and left overnight at 37 °C in a humid chamber. After covalent binding, the other coverslip was removed, PAA gels were activated and ECM-treated as described above. Monolayers were cultured as described above with additional 12.5µg/ml Metronidazole (Braun) and 4µg/ml Ciprofloxacin (Panpharma) in the culture medium and stretched for 30sec for 24h, 10% strain, 0.26 Hz.

### Generation of Lgr5-DTR-EGFP/K-GECO1 organoid line

The pLV-K-GECO1-IRES-Puro plasmid was a kind gift from Hugo Snippert (University Medical Center Utrecht, The Netherlands), generated through In-Fusion cloning of the insert of the original pcDNA-K-GECO1 vector (Addgene plasmid #105865)(*33*) into a lentiviral pLV vector. Lgr5-DTR-EGFP ileum organoids with stable expression of the K-GECO1 calcium sensor were generated by transducing primary organoids derived from the ileum of Lgr5-DTR-EGFP mice (*60*) with lentivirus containing the pLV-K-GECO1-IRES-Puro plasmid and selection based on Puromycin resistance.

### Generation of 7TG organoid line

Lentivirus production and *R26^mTom/mGFP^* organoids transduction have been extensively described elsewhere (*57*). Lenti-7TG (7xTcf-eGFP) plasmid (Addgene plasmid# 24314), pMD2.G (Addgene plasmid# 12259) and psPAX2 (Addgene plasmid# 12260) were used for lentiviral production.

### Live imaging of intracellular calcium

For calcium imaging experiments, Lgr5-DTR-EGFP/K-GECO1 ileum organoids were seeded as monolayers as described above. Imaging was performed 4 days after seeding once mature crypt-like domains were established. Time-lapse K-GECO1 images were acquired with 5 seconds time interval on a Nikon Spinning Disc confocal microscope (Yokogawa CSU-W1) using a 40X water objective (NA = 1.15) in a temperature- and CO_2_-controlled incubator, using NIS-Elements software. K-GECO1 time-lapse movies are presented as 6 µm MAX Z-projections and processed using ImageJ software.

For calcium analyses, K-GECO1 fluorescence intensity was measured in max Z-projections over time within segmented Lgr5-eGFP^+^ regions of crypt-like domains. Of individual Lgr5-eGFP^+^ regions the K-GECO1 response to Yoda1 stimulation was determined by calculating (F- F_0_)/F_0,_ with F being peak fluorescence intensity and F_0_ the average intensity of the 10% lowest values before Yoda1 addition.

### Single cell RNA-seq and analysis

As previously described (*55*), crypts were isolated from mouse small intestine using 20mM EDTA/PBS1X (Gibco) for 25min at 4°C with continuous shaking. Intestines pieces were then incubated in 1.5% sucrose/1% sorbitol/PBS1X solution and crypts were dissociated by vigorous shaking. Following 70um filtering and centrifugation, crypts were incubated with TrypLE 1X at 37°C for 5min. After pipetting multiple times until the single-cell suspension was attained, the solution is passed through a 40um strainer and centrifugated. Trypan blue staining of the single-cell suspension was used to confirm >80% viability. An estimated 10000 cells were loaded for 10x Genomics single-cell isolation and library preparation was performed according to manufacturer’s recommendations. Illumina Hiseq 3000 was used to sequence the sample. The whole intestine of 3 control or PIEZO mutant mice were pooled per dataset. Sequences from scRNA-seq were processed with Cellranger (v.2.0.0) software (10x Genomics). In short, demultiplexing, UMI (unique molecular identifier) collapsing and alignment to the mm10 mouse transcriptome was performed. The raw data for control mice generated by Cellranger were then read into the Seurat R package (*61*) with at least 200 genes present in each cell and at least one cell. Quality control and normalization were then performed. For each cell, quality control metrics were calculated, including the total number of counts, the proportion of counts in mitochondrial genes, and the number of genes expressed by cell. Specifically, genes with non-zero counts in at least three cells were retained to remove any low-abundance genes that might interfere with statistical analysis. The gene expression was log-normalized, then scaled. A cell cycle score was calculated based on expression of various cell-cycle associated genes, pre-defined within Seurat. This score was treated as a regression variable upon data scaling to prevent cells from clustering together based on cell cycle phase. Principal components were then calculated on the basis of the 2000 most variable genes in the dataset (excluding those associated with the cell cycle) and a scree plot was used to determine the top principal components. Cells were then divided into unsupervised clusters using a shared-nearest-neighbour modularity optimization-based clustering algorithm (Smart local moving algorithm (*62*)) and then plotted using the UMAP method (*63*). Cells were initially clustered at low resolution and immune cells were removed using *Ptprc* as a marker. Epithelial cells were re-clustered at high resolution and expressed genes for each cluster were identified using the Wilcox test in Seurat and the top genes (<0.05 P value and >0.25 log fold change) were used to identify different cell types. To test how single-cell clusters identified in control intestinal epithelia were altered upon *Piezo* deletion, scRNA-seq dataset from *Piezo* mutant was first processed as above by using the same parameters that were applied to process the control dataset. Next, FindTransferAnchors function within Seurat was used to compare the two datasets, where the control clusters were used as a reference map to infer clusters in the mutant dataset. A UMAP plot of the *Piezo* mutant cells mapped back to clusters calculated from the control data was then generated. All data plotting and analyses were done in R.

### Atomic Force Microscopy

The small intestine of a B6N mouse was dissected and flushed gently with cold 1X PBS to remove fecal content. The jejunum was cut into segments of a few centimeters, filled with OCT using a syringe, placed in a cryomold, and embedded in OCT. The embedded tissue was fresh frozen in a bath of 100% EtOH, cooled down with dry ice and then stored at −80°C. Using a cryostat (CM3050 S, Leica) the OCT blocks were cut at −20 °C chamber temperature and −15 °C object temperature to generate 30 µm-thick transversal sections. The sections were transferred onto FluoroDish glass bottom dishes (FD35-100, wpi) precoated with 0.28 mg/ml Cell-Tak (Corning, 354240) overnight at 37°C, followed by one wash with water and air drying. The glass bottom dishes containing tissues were stored at −80°C for up to 2 weeks. On the day of the measurements, the sections were thawed and immediately permeabilized and blocked in a buffer containing 0.5% Triton X-100 (Sigma) and 10% goat serum (Gibco) for 15 min at RT. The tissues were then incubated with anti-laminin alpha 2 (1:100, Sigma, L0663) for 30 min at RT, washed 2 x 2 min with 1X PBS, and incubated with anti-rat 546 (1:500, Thermo Fischer) and DAPI (1:10000, Life Technologies) for 30 min at RT. For the measurements, the tissues were immersed in PBS supplemented with a protease inhibitor cocktail (Roche, 11873580001) and Pen/Strep (Gibco).

An MLCT-O10 probe (Bruker) was mounted on a CellHesion 200 AFM (Bruker) which is integrated into an Eclipse Ti inverted light microscope (Nikon). The sensitivity and the spring constant of the cantilever at position D of the probe were calibrated in air via contact-based calibration and thermal oscillation (spring constant 0.4 to 0.5 N/m) and then a 10 µm-diameter glass bead was glued to the tip of the cantilever. Before the measurements, the cantilever was coated with 1 % Pluronic F-127 (Sigma, P2443) for 1 h at 25°C, rinsed in PBS and the sensitivity was re-calibrated in liquid. The laminin alpha 2 staining was used to identify the basement membrane (BM) to be probed. For each crypt top and bottom, measurements were performed over three consecutive 20 µm-long horizontal lines, spaced by 2 µm, centered on the BM and oriented perpendicular to the BM (see Fig. 5G). The measurements were done every 0.5 µm (i.e. 41 points per line). If necessary, the sample was rotated to position the BM vertically. For the crypt top, care was taken to avoid probing two BMs. Measurements were run at 25°C as follows: the samples were approached with a speed of 5 µm/s and indented with 0.4 µm/s speed up to a maximum force of 6 nN. Resulting force-time curves were analyzed using the JPK Data Processing Software using the Hertz-model with a fitting range of −0.15 to 1 µm. Force-time curves without a clear contact point or with noise were discarded. Upon data visualization, datasets for which no matching peak was identified among the 3 lines were excluded from the peak value analysis (Table in Fig. S7J). An exemplary selected data set is represented in Fig. 6H. For all selected data sets, the peaks of each line were aligned manually (Fig. S7K) and the average of these 3 peak values was used to determine the Young’s modulus of the crypt.

### Quantification and statistical analysis

No statistical methods were used to predetermine sample size. The investigators were blinded to allocation during experiments and outcome assessment. No animal has been excluded from analysis and no randomization method has been applied in this study. Every experiment has been performed at least twice with the same outcome. The number of independent experimental replications, the definition of centre, variation (mean ± SD) and statistical test (*P* value) were reported in each corresponding figure legend. All statistical analyses were performed with GraphPad Prism software.

## ACKNOWLEDGEMENTS

The authors would like to thank the Cell and Tissue Imaging Platform and the CurieCoreTech cytometry facility of Institut Curie. We thank the members of the Vignjevic lab (Institut Curie) for their advice and comments during this study. We thank the Fre lab for their technical help with the 3D organoids.

## Authors contributions

M.B.B. conceived and designed the project and performed most of the experiments, analysed the data, and wrote the manuscript; A.R-B. initiated the project; R.H. performed the calcium imaging experiments; M.W. analysed the scRNA-seq dataset; G.G. and S.D. designed and developed the cell stretching device; R.B helped with the 3D organoid lines. A.K.W., A.Af., X.C. and X.H. helped with the mice models; M.G. supervised the calcium imaging experiments; J.L.W. supervised scRNAseq experiments and analysis; A.A. performed and analysed scRNA-seq; L.P, L.D., M.B. and A.D.M performed and analyzed the AFM experiments, T.-H.K. and D.M.V. supervised the project, and revised the manuscript. All authors read and agreed on the manuscript.

## Data and materials availability

This work analyzes a publicly available dataset: GEO accession number GSE143915 (Fig. 1A). The scRNAseq raw sequencing data generated in this study will be provided upon request. All data are available in the main text or the supplementary materials.

## SUPPLEMENTARY MATERIALS

Supplementary Figures S1 to S7; Table S1 and S2; Movies S1 to S7.

