## supplementary txt for "PIEZO-dependent mechano-sensing of the niche is essential for intestinal stem cell fate decision and maintenance"

[1]Institut Curie, PSL Research University, CNRS UMR 144, F-75005 Paris, France; [2]Center for Molecular Medicine, University Medical Center Utrecht and Utrecht University, Utrecht, the Netherlands; [3]Program in Developmental & Stem Cell Biology, The Hospital for Sick Children, Toronto, Ontario M5G 0A4, Canada; [4]Department of Molecular Genetics, University of Toronto, Toronto, Ontario M5S 1A8, Canada; [5]Department of Biological Sciences, Faculty of Science, University of Calgary, Calgary, Alberta, Canada; [6]Arnie Charbonneau Cancer Institute, Cumming School of Medicine, University of Calgary, Calgary, Alberta, Canada; [7]Cell Biology and Biophysics Unit, European Molecular Biology Laboratory, 69117, Heidelberg, Germany ; [8]Institut Curie, IPGG, PSL Research University, CNRS UMR 168, F-75005 Paris, France; [9]Arthur and Sonia Labatt Brain Tumour Research Centre, The Hospital for Sick Children, Toronto, Ontario, Canada.[10]Centre for Systems Biology, Lunenfeld-Tanenbaum Research Institute, Mount Sinai Hospital, Toronto, Ontario, Canada; [11]Department of Laboratory Medicine, St. Michael's Hospital, Toronto, Ontario M5B 1W8, Canada.

*__Corresponding authors:__

Meryem B. Baghdadi,
Tae-Hee Kim,
Danijela Matic Vignjevic,

### These authors contributed equally to this work

#### SUPPLEMENTARY FIGURES

**Figure S1: Control smFISH**

**(A)** Representative images of smFISH for negative and positive control probes. DAPI stains DNA.

**Figure S2: *Piezo2* compensates for *Piezo1* loss to maintain intestinal homeostasis.**

**(A)** Experimental scheme of WT control (Ctr) and *Piezo1* (*Piezo1$^{cKO}$; Villin-creERT2; Piezo1$^{Flox}$*) or *Piezo2* (*Piezo2$^{cKO}$; Villin-creERT2; Piezo2$^{Flox}$*) conditional knock-out mice. Mice were injected intraperitoneally (ip) with tamoxifen once a day for 5 consecutive days and sacrificed at day 6 (D6). **(B)** smFISH for *Piezo1* expression in WT control (Ctr) and *Piezo1* conditional KO mice (*Piezo1$^{cKO}$*). Quantification of the number of *Piezo1* transcripts in Ctr and *Piezo1$^{cKO}$* shows efficient recombination upon tamoxifen injections in mutant mice (n=50 crypt/villus unit from N=4 mice). **(C)** smFISH for *Piezo2* expression in WT control (Ctr) and *Piezo2* conditional KO mice (*Piezo2$^{cKO}$*). Quantification of the number of *Piezo2* transcripts in Ctr and *Piezo2$^{cKO}$* shows efficient recombination upon tamoxifen injections in mutant mice (n=50 crypt/villus unit from n=4 mice). **(D)** Representative images of Hematoxylin/Eosin (H&E) staining of intestinal transverse sections from control (Ctr), *Piezo1* (*Piezo1$^{cKO}$*) and *Piezo2* (*Piezo2$^{cKO}$*) conditional KO mice. Quantification of crypt length (n=150 crypts from N=3 mice) in Ctr, *Piezo1$^{cKO}$* and *Piezo2$^{cKO}$* mice. **(E)** Representative images of Alcian Blue staining of intestinal transverse sections and quantification of secretory cell numbers (Alcian Blue-positive) in Ctr, *Piezo1$^{cKO}$* and *Piezo2$^{cKO}$* mice (n=150 crypts from N=3 mice). **(F)** Immunostaining for OLFM4 and ISC quantification in Ctr, *Piezo1$^{cKO}$* and *Piezo2$^{cKO}$* mice. **(G)** smFISH for *Piezo2* expression in intestinal transversal sections of WT control (Ctr) and *Piezo1* conditional KO mice (*Piezo1$^{cKO}$*; *Villin-creERT2;Piezo1$^{Flox}$*). E-cadherin immunostaining shows the epithelium (E-CAD, white) and DNA (DAPI, blue). Insets show increase of *Piezo2* expression in the crypt region (n=150 crypt/villus unit from N=4 mice). Scale bar, 50µm and 10µm in insets. Error bars indicate SD. Two-tailed unpaired Student's *t*-test (D-F), Kolmogorov-Smirnov test (B, C, G) and *P* values are shown in panels. n.s: non-significant. Box edges show 25$^{th}$ and 75$^{th}$ percentiles, central point is median, and error bars represent minimum and maximum values.

**Figure S3: Characterization of *Piezo1;Piezo2* conditional double knock-out mice.**

Representative images of smFISH for **(A)** *Piezo1* in control (Ctr; *Cre-; Piezo1$^{Flox}$;Piezo2$^{Flox}$*) and *Piezo1* and *Piezo2* double conditional knock-out mice (*Piezo$^{dbKO}$*; *Villin-creERT2; Piezo1$^{Flox}$;Piezo2$^{Flox}$*). Quantification of the number of *Piezo1* transcripts in Ctr and *Piezo$^{dbKO}$*

showing efficient recombination upon tamoxifen injections in double mutant mice (n=50 crypt/villus unit from N=4 mice). Insets show absence of *Piezo1* transcripts in intestinal crypts while still present in the stroma (arrow). E-Cadherin (ECAD) immunostaining is used to visualize intestinal epithelial cells and DAPI (DNA). **(B)** PIEZO immunostaining on Ctr and *Piezo$^{dbKO}$* intestines validating PIEZO1 antibody used in Fig. 1C. Scale bar, 25µm. **(C)** *Piezo2* transcripts in Ctr and *Piezo$^{dbKO}$* showing efficient recombination upon tamoxifen injections in double mutant mice (n=50 crypt/villus unit from N=4 mice). Insets show absence of *Piezo2* transcripts in intestinal crypts while still present in the stroma (arrow). E-Cadherin (ECAD) immunostaining is used to visualize intestinal epithelial cells and DAPI (DNA). **(D)** Representative image of smFISH for *Lgr5* transcript in crypts of Ctr and *Piezo$^{dbKO}$* mice with E-Cadherin (ECAD) immunostaining to visualize intestinal epithelial cells and DAPI (DNA). Quantification of *Lgr5* transcripts in control and double mutant mice. Scale bar, 10µm. **(E)** EdU pulse (24h) and labelling in Ctr and *Piezo$^{dbKO}$* mice. Quantification of the EdU front was defined as the distance measured from the villus bottom to the front-most EdU-labelled epithelial cell (n=3 mice). Scale bar, 50µm. **(F)** TUNEL staining in intestinal transverse sections from control and *Piezo$^{dbKO}$* mice four days post-tamoxifen induction. Scale bar, 50µm and 10µm in insets. **(G)** *Piezo1* and *Piezo2* transcript analysis (qRT-PCR) shows recombination efficiency in 3D organoids generated from Ctr and *Piezo$^{dbKO}$* mice and treated with 4-hydroxytamoxifen (4'OHT) for 48h *in vitro* (n=3 independent experiments). **(H)** smFISH of *Fabp* (absorptive marker), *Tff3* (secretory marker) and *Lgr5* (stem marker) on 3D organoids from *Piezo$^{dbKO}$* mice and induced or not with 4'OHT. Scale bar, 50µm and 10µm in insets. Error bars indicate SD. Kolmogorov-Smirnov test (A, C, D), Two-tailed unpaired Student's *t*-test (E-G), and *P* values are shown in panels. Box edges show 25th and 75th percentiles, central point is median, and error bars represent minimum and maximum values.

**Figure S4: scRNAseq of intestinal crypt cells isolated from *Piezo$^{dbKO}$* mice.**

**(A)** Differential gene expression (DGE) analysis shows differentially expressed transcription factors in mouse intestinal epithelial cells and cluster cell identity. EC: Enterocytes. **(B)** UMAP plot of stem cell markers (*Lgr5, Ascl2)* expression in control (Ctr; *Cre-; Piezo1$^{Flox}$;Piezo2$^{Flox}$*) and *Piezo1* and *Piezo2* double conditional knock-out (*Piezo$^{dbKO}$*; *Villin-creERT2; Piezo1$^{Flox}$;Piezo2$^{Flox}$*). **(C)** UMAP plot of proliferation markers (*Mcm6*, *Mcm5*, *Pcna)* expression in Ctr and *Piezo$^{dbKO}$* mice. **(D)** Immunostaining of OLFM4 (stem), LYZ (Paneth cells), MUC2 (Goblet cells), ChgA (enteroendocrine cells), DCLK (Tuft cells), ANPEP (enterocytes) and EdU labeling (proliferative/TAs) labeling in Ctr and *Piezo$^{dbKO}$* mice 4 days post-tamoxifen induction. Matching quantifications are presented below each representative images. (n=50 crypts from 4 mice). Scale bar, 50µm. Error bars indicate SD. Two-tailed unpaired Student's *t*-test (D), *P* values are shown in panels, error bars represent minimum and maximum values.

**Figure S5: Paneth cells alterations are not responsible for ISC defect upon *Piezo* deletion**

**(A)** Heatmap of all differentially expressed genes in Paneth cell cluster (#12) in Ctr and *Piezo*$^{dbKO}$ mutant mice. **(B)** Heatmap of NOTCH ligand gene expression in Paneth cell cluster (#12) in Ctr and *Piezo*$^{dbKO}$ mutant mice. **(C)** Heatmap of WNT ligand gene expression in Paneth cell cluster (#12) in Ctr and *Piezo*$^{dbKO}$ mutant mice (left). Violin plot showing *Wnt3* expression level in Ctr and KO Paneth cells (right).

**Figure S6: PIEZO inhibition alters WNT signaling**

**(A)** Lentiviral transduction of 7TG plasmid in *R26*$^{mT/mG}$ organoids showed 30% of transduction efficiency. This representative image shows the GFP signal intensity (FIRE LUT) in 7TG-positive organoid compared to 7TG-negative one. **(B)** Representative images of brightfield time-lapse imaging of control (Ctr), $GdCl_3$, and GsMTx4-treated organoids. See Movie S5. Quantification of *de novo* crypts formed per organoid (n=100 organoids from n=3 independent experiments). Scale bar, 50µm. Error bars indicate SD. Two-tailed paired Student's *t*-test (B). *P* values are shown in panels and error bars represent minimum and maximum values.

**Figure S7: Characterization and validation of the 2D monolayer model**

**(A)** Experimental scheme of generation of monolayer from crypt organoids isolated from non-induced *Piezo1* and *Piezo2* double conditional knock-out mice (*Piezo*$^{dbKO}$; *Villin-creERT2; Piezo1*$^{Flox}$*;Piezo2*$^{Flox}$). After the monolayers are formed (3 to 5 days post-seeding), 4-hydroxytamoxifen (4'OHT) is added to the culture medium for 48h to induce recombination. **(B)** *Piezo1* and *Piezo2* transcript analysis shows recombination efficiency in 2D organoid monolayers generated from Ctr and *Piezo*$^{dbKO}$ mice and treated with 4'OHT for 48h *in vitro* (n= 4 monolayers). **(C)** OLFM4 immunostaining in monolayers from *Piezo*$^{dbKO}$ treated with 4'OHT. Quantification of percentage of crypt-containing stem cells (OLFM4-positive) shows stem cell loss upon *Piezo1* and *Piezo2* deletion. ECAD immunostaining shows epithelial cells (n= 25 crypts from N=3 independent experiments). **(D)** LYSOZYME (LYZ) immunostaining in monolayer from *Piezo*$^{dbKO}$ treated with 4'OHT. Quantification of percentage of crypt-containing Paneth cells (LYZ-positive) shows Paneth cell loss upon *Piezo1* and *Piezo2* deletion. ECAD immunostaining shows epithelial cells (n= 25 crypts from N=3 independent experiments). **(E)** EdU labelling and quantification in monolayers from *Piezo*$^{dbKO}$ treated with 4'OHT (n= 35 crypts from N=3 independent experiments). **(F)** UEA labeling (all secretory cells) and quantification in monolayers from *Piezo*$^{dbKO}$ treated with 4'OHT. (n= 30 crypts from N=3 independent experiments). Scale bar, 10µm. **(G)** EZRIN immunostaining showing absorptive cells in monolayers from *Piezo*$^{dbKO}$ treated with 4'OHT. **(H)** Baseline of $Ca^{2+}$ transients (K-GECO) in Lgr5-GFP+ crypts of monolayers seeded on 1.5KPa and 18kPa substrates without any stimulation i.e. before addition of Yoda1. (n=50 crypts from N=3 independent experiments). See Movies S6 and S7. **(I)** Representative images of OLFM4 and membrane tdTomato (Memb) immunostaining on monolayer cultured on substrate of 5kPa, 11kPa and 30 kPa, treated with PIEZO inhibitor (GsMTx4) or activator (Yoda1). Quantification of the stem cell area over non-treated control (Ctr) (n= 30 crypts from N=3 independent experiments). Scale bar, 25µm. **(J)** Young's Modulus (kPa) as the average of all values (raw data) for all bottom and top crypt datasets (corresponding to Fig. 5I left graph); or as the average of peak values for selected bottom and top crypt datasets (corresponding to Fig. 5I right graph). **(K)** Young's Modulus traces for the 3 lines represented in Fig. 5H before (top) and after (bottom) alignment. Highlighted measurement corresponds to the crypt presented in Fig. 5G. Scale bar, 25µm (C, D, F, G). Error bars indicate SD. Two-tailed paired Student's *t*-test (B-D, F), Kolmogorov-Smirnov test (E, H) and *P* values are shown in panels; *n.s*: non-significant. Box edges show 25th and 75th percentiles, central point is median, and error bars represent minimum and maximum values.

**Figure S8: PIEZO activation permeates calcium in 3D organoids**

**A)** Time series and corresponding traces of calcium influx (FIRE LUT) in crypts from Lgr5-K-GEKO organoid before (-) and after addition (+) of the PIEZO activator, Yoda1 or DMSO (control vehicle). Each trace represents the average of two individual Lgr5-GFP+ crypts within a single organoid (n=6 crypts/condition from 3 individual organoids are depicted per graph). Scale bar, 25µm. **(B)** Max intensity increase of calcium ($dF_{max}/F_0$) after Yoda1 or DMSO in individual Lgr5-GFP+ crypts (n=5-8 crypts/condition in each experiment, from N=3 independent experiments). Two-tailed paired Student's *t*-test. *P* values are shown in panel. Box edges show 25th and 75th percentiles, central point is median, and error bars represent minimum and maximum values.

**Supplementary table 1: list of RT-qPCR primers used in this study.**

| Gene | Forward | Reverse |
|---|---|---|
| *Tbp* | ATCCCAAGCGATTTGCTG | CCTGTGCACACCATTTTTCC |
| *Rpl13* | GTGGTCCCTGCTGCTCTCAAG | ATAGTGCATCTTGGCCTTTT |
| *Lgr5* | GACAATGCTCTCACAGAC | GGAGTGGATTCTATTATTATGG |
| *Olfm4* | GCCACTTTCCAATTTCAC | GAGCCTCTTCTCATACAC |
| *Axin2* | GGACTGGGGAGCCTAAAGGT | AAGGAGGGACTCCATCTACGC |
| *Lrig* | TTCCTTACCGGTGAGACTGG | CCATCACTGTGCCAACACTT |
| *Lyz1a* | GGAATGGATGGCTACCGTGG | CATGCCACCCATGCTCGAAT |
| *Defa5* | TATCTCCTTTGGAGGCCAAG | TTTCTGCAGGTCCCAAAAAC |
| *Ki67* | ATCATTGACCGCTCCTTTAGGT | GCTCGCCTTGATGGTTCCT |
| *Piezo1* | AGGACTTCCCCACCTATTGG | CCAGGGATGAGGATACTGGAAAA |
| *Piezo2* | AGAGTCGGAAAAGAGATACCCTC | CCAGACGATACAGATGAGAAGGA |

**Supplementary table 2: Volumes of reagents to prepare different stiffness PAA gels (all volumes are in µl).**

| Stiffness (Young Modulus) | PBS | Acrylamide 40% | Bis-acrylamide 2% | APS | TEMED |
|---|---|---|---|---|---|
| **1.5 kPa** | 407,5 | 68,75 | 11 | 2,5 | 0,25 |
| **5 kPa** | 382,95 | 93,3 | 11 | 2,5 | 0,25 |
| **11 kPa** | 368,5 | 93,75 | 25 | 2,5 | 0,25 |
| **18 kPa** | 352,85 | 94,4 | 40 | 2,5 | 0,25 |
| **30 kPa** | 299,75 | 150 | 37,5 | 2,5 | 0,25 |

**Supplementary movies:**

**Movie S1:** Videomicroscopy of control crypt organoid growth. Images were taken every hour for 36h.

**Movie S2:** Videomicroscopy of crypt organoid treated with PIEZO inhibitor $GdCl_3$. Images were taken every hour for 36h.

**Movie S3:** Videomicroscopy of control crypt treated with PIEZO inhibitor GsMTx4. Images were taken every hour for 36h.

**Movie S4:** Videomicroscopy of control crypt treated with PIEZO activator Yoda1. Images were taken every hour for 36h.

**Movie S5:** Videomicroscopy of Lgr5-GFP;K-GECO1 crypt from monolayer cultured on 1.5kPa substrate before and after addition of Yoda1.

**Movie S6:** Videomicroscopy of Lgr5-GFP;K-GECO1 crypt from monolayer cultured on 18kPa substrate before and after addition of Yoda1.

**Movie S7:** Videomicroscopy of Lgr5-GFP;K-GECO1 3D organoid before and after addition of DMSO (vehicle control).

**Movie S8:** Videomicroscopy of Lgr5-GFP;K-GECO1 3D organoid before and after addition of Yoda1.
