## supplementary figures for "PIEZO-dependent mechano-sensing of the niche is essential for intestinal stem cell fate decision and maintenance"

A

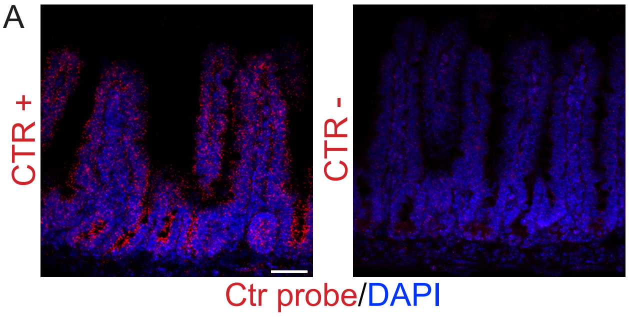

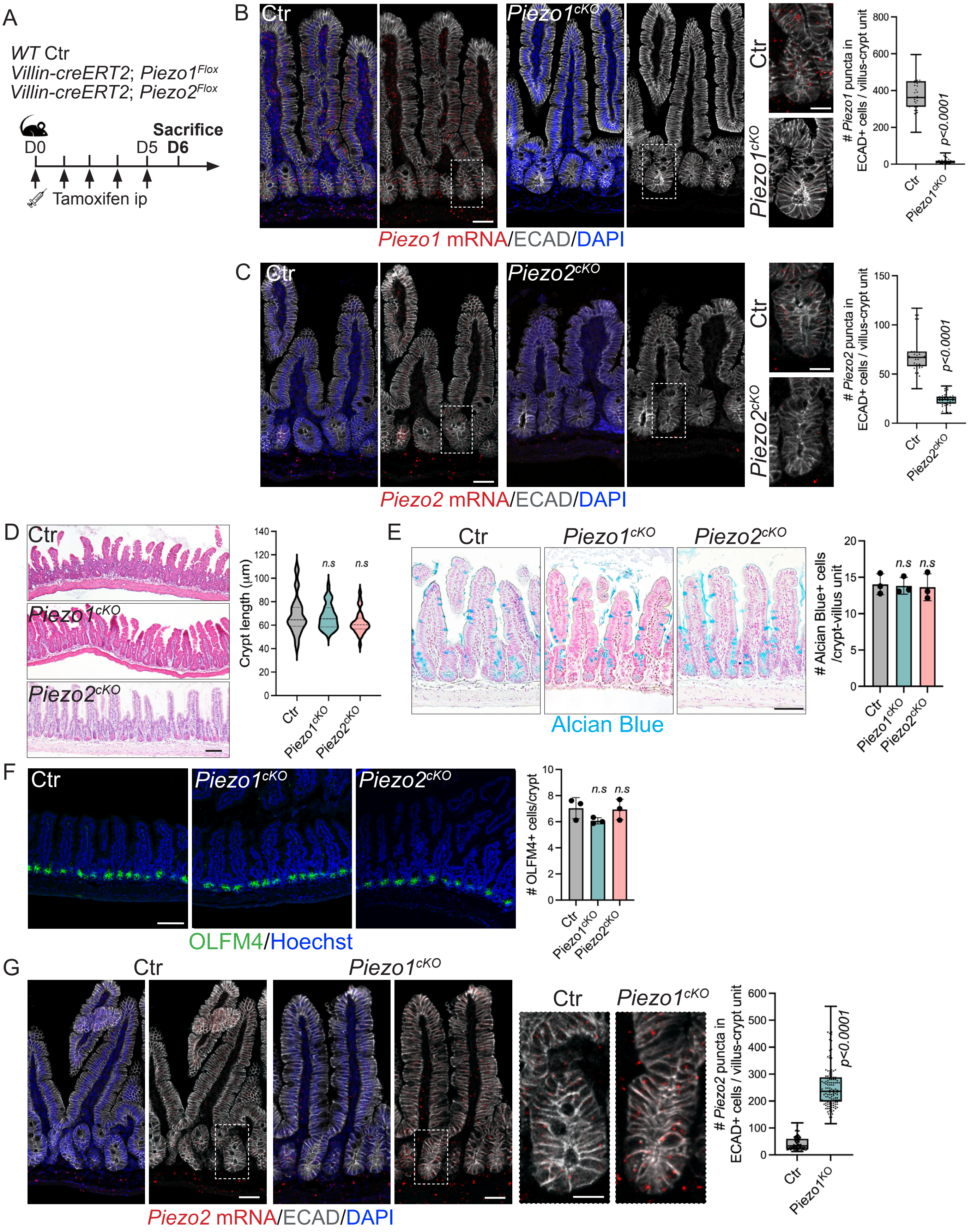
A
WT Ctr
Villin-creERT2; Piezo1Flox
Villin-creERT2; Piezo2Flox
Sacrifice
D0
D5 D6
Tamoxifen ip
B
Ctr
Piezo1cKO
Piezo1 mRNA/ECAD/DAPI
### Piezo1 puncta in ECAD+ cells / villus-crypt unit
p<0.0001
C
Ctr
Piezo2cKO
Piezo2 mRNA/ECAD/DAPI
### Piezo2 puncta in ECAD+ cells / villus-crypt unit
p<0.0001
D
Crypt length (µm)
n.s
E
Alcian Blue
### Alcian Blue+ cells /crypt-villus unit
F
OLFM4/Hoechst
### OLFM4+ cells/crypt
G
Piezo2 mRNA/ECAD/DAPI
### Piezo2 puncta in ECAD+ cells / villus-crypt unit
Piezo1KO

Baghdadi_FIGURE S3

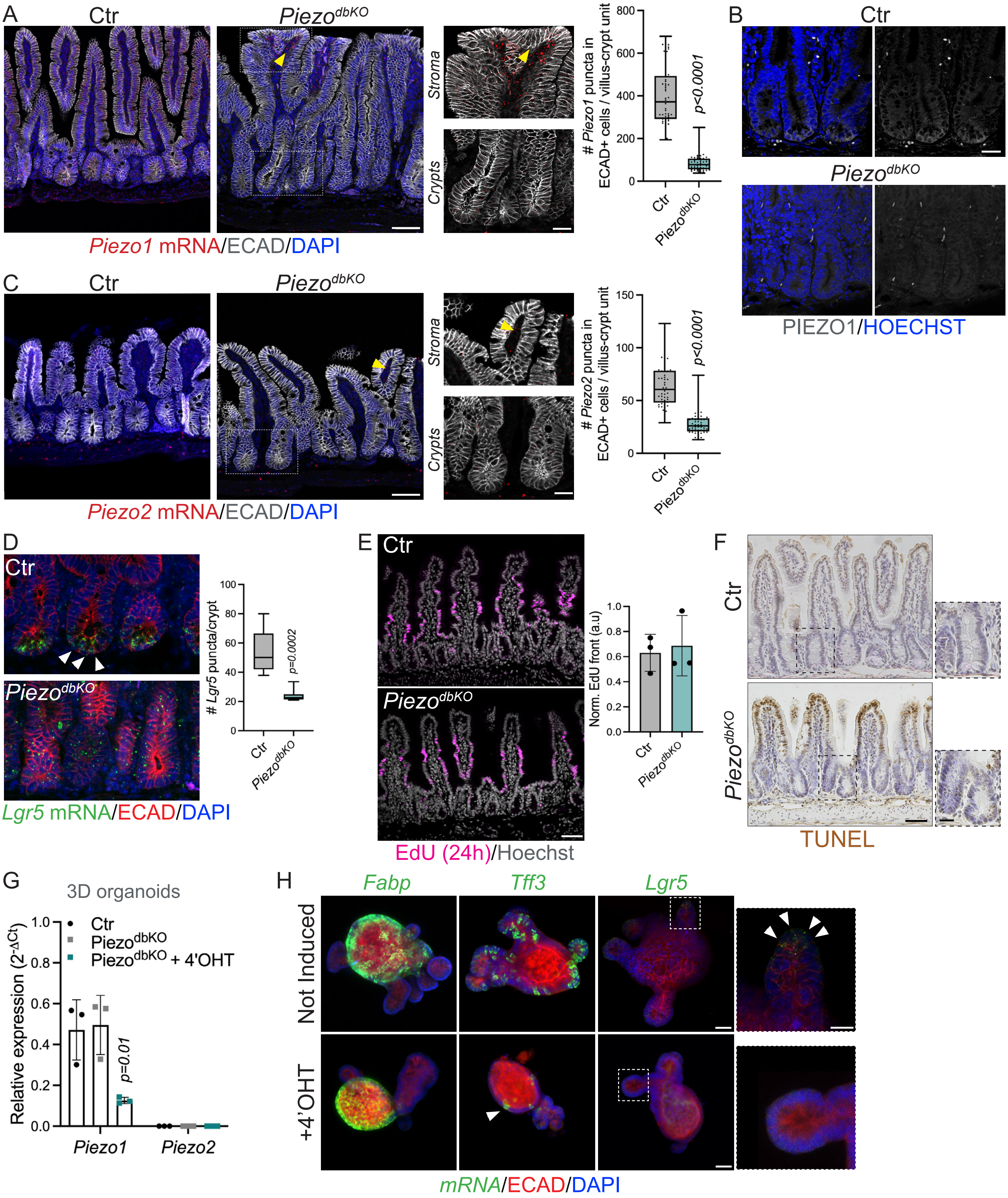

Baghdadi_FIGURE S4

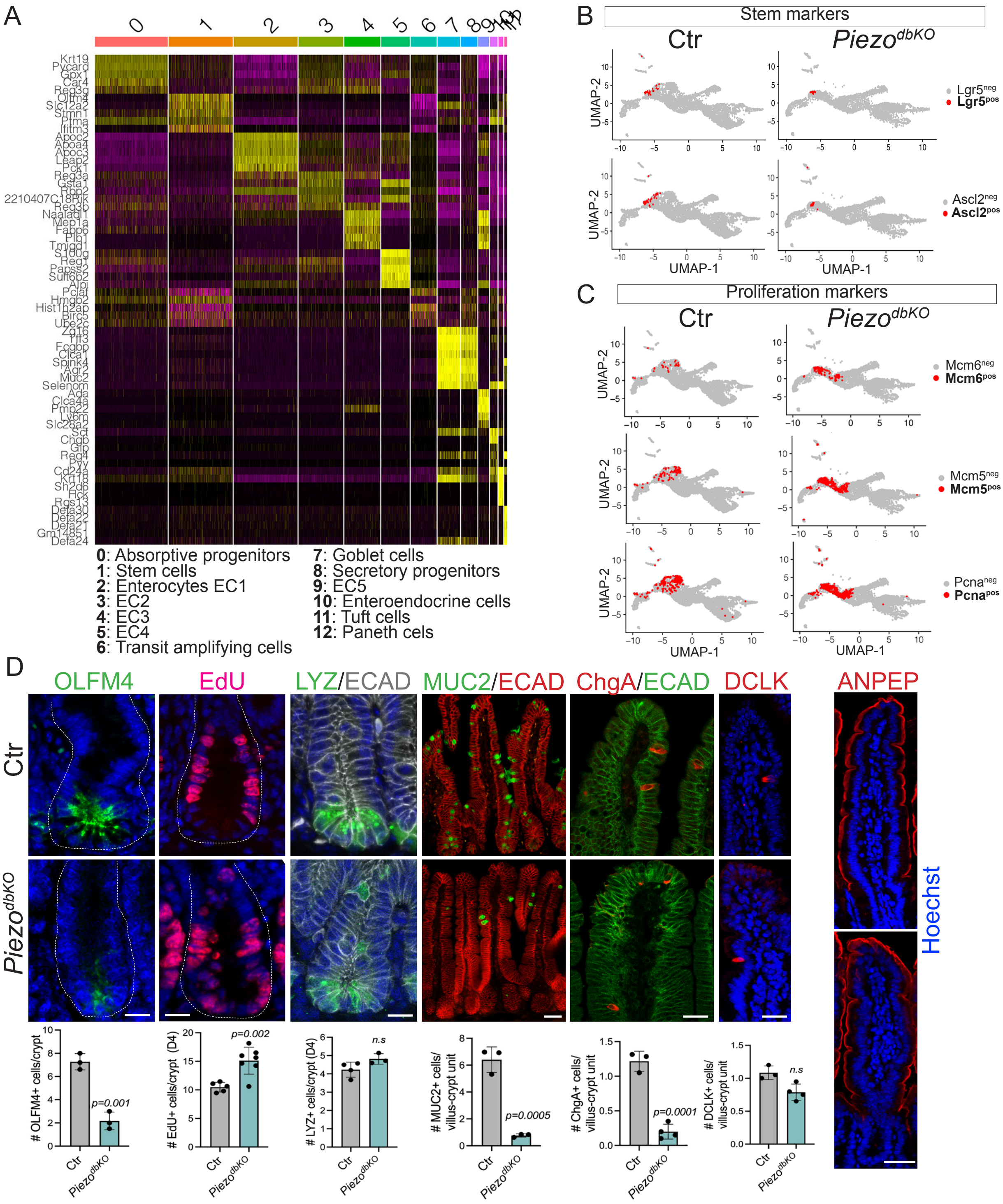

Baghdadi_FIGURE S5

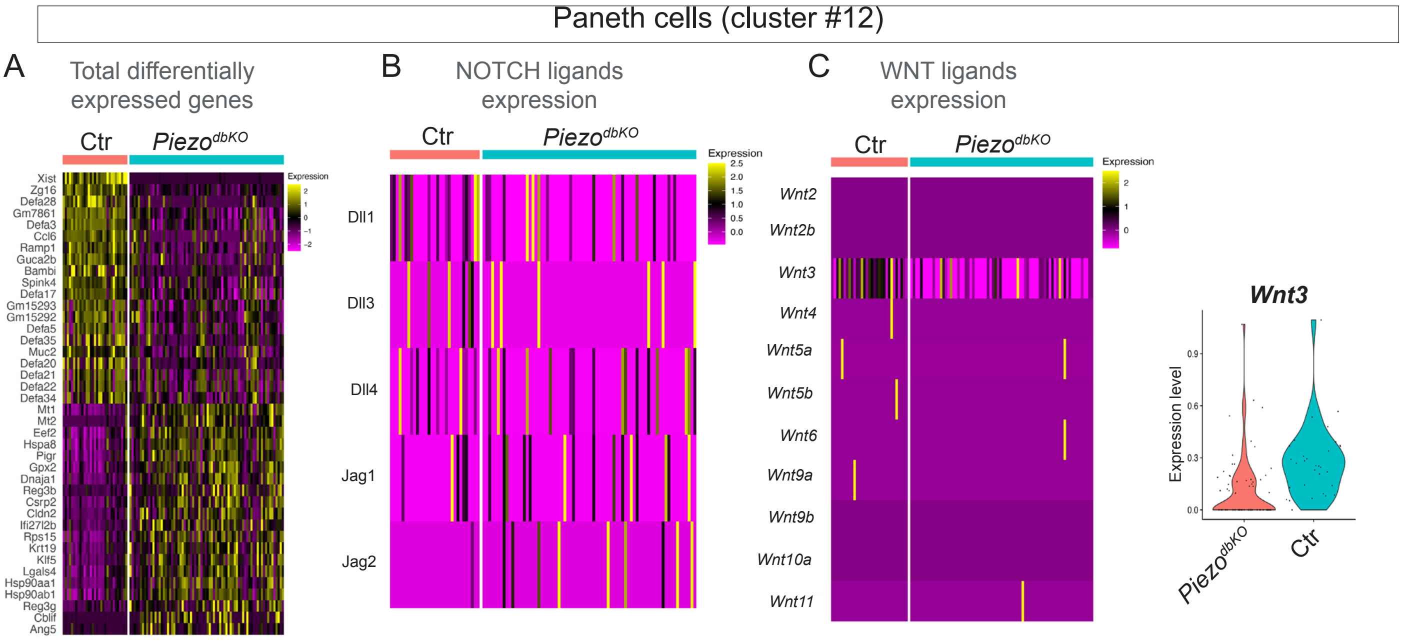

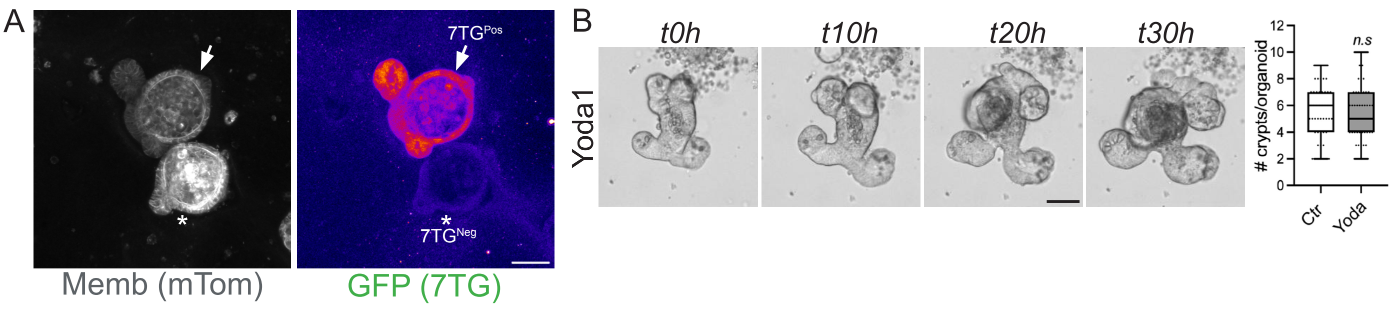
A
7TGPos
7TGNeg
Memb (mTom)
GFP (7TG)
B
t0h
t10h
t20h
t30h
Yoda1
### crypts/organoid
n.s
0
2
4
6
8
10
12
Ctr
Yoda

### Baghdadi_FIGURE S7

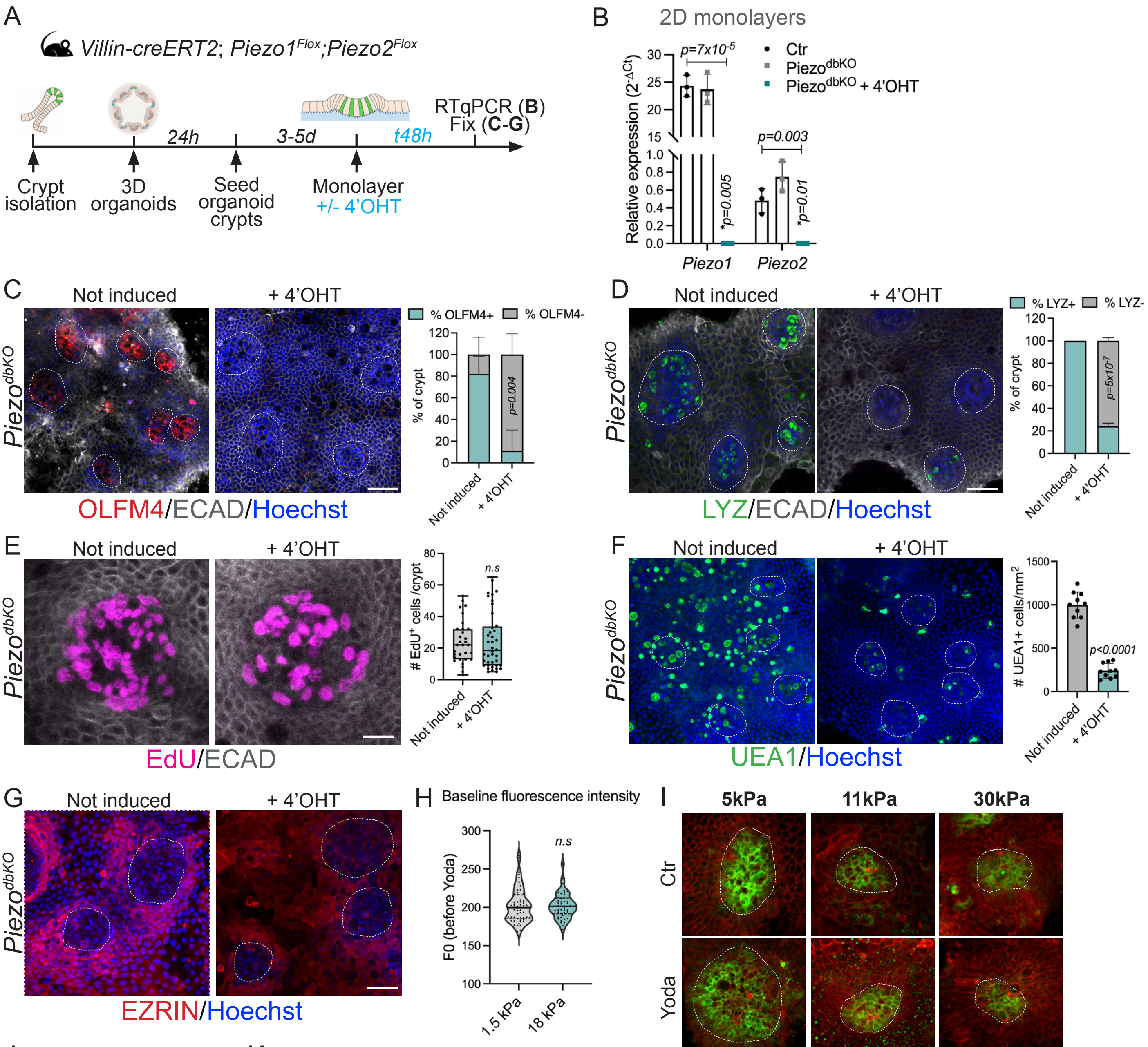


J

| | Average Young's modulus (kPa) | | Peak Young's modulus (kPa) | |
|---|---|---|---|---|
| | Bottom | Top | Bottom | Top |
| Crypt 1 | 2,00 | n.a. | 7,24 | n.a. |
| Crypt 2 | 2,96 | n.a. | 19,77 | n.a. |
| Crypt 3 | 4,93 | 1,43 | 21,89 | 5,48 |
| Crypt 4 | 3,41 | 1,03 | 4,73 | 4,46 |
| Crypt 5 | 2,93 | 1,08 | 20,93 | 3,91 |
| Crypt 6 | 2,38 | 0,95 | 12,12 | n.a. |
| Crypt 7 | n.a. | 1,17 | n.a. | 3,62 |
| Crypt 8 | n.a. | 0,74 | n.a. | 3,06 |
| Crypt 9 | n.a. | 2,00 | n.a. | 5,25 |
| Crypt 10 | 1,41 | 1,20 | 6,77 | 4,47 |
| Crypt 11 | 0,51 | 0,56 | 1,90 | 2,21 |
| Crypt 12 | 1,78 | n.a. | | |
| Crypt 13 | 3,54 | 0,28 | | |
| Crypt 14 | 8,78 | n.a. | | |
| Crypt 15 | 3,35 | n.a. | | |
| **Average** | **3,17** | **1,04** | **11,92** | **4,06** |

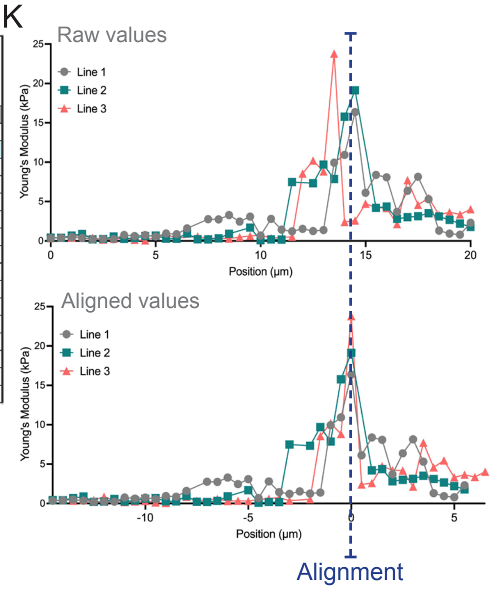


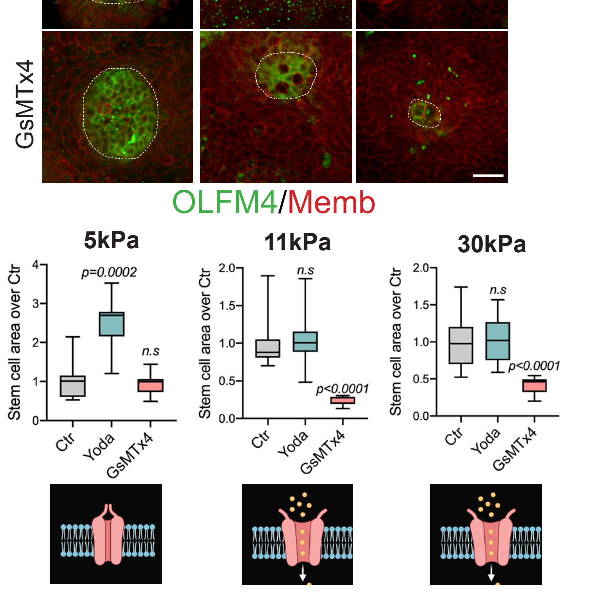

Baghdadi_FIGURE S8

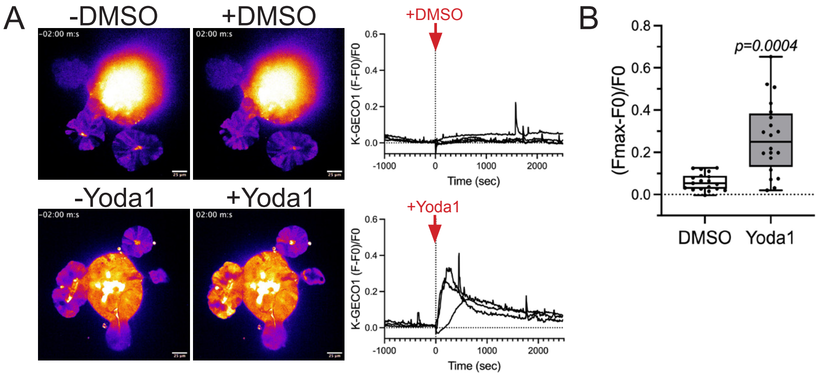
